# An Autoimmune Transcriptional Circuit Driving Foxp3^+^ Regulatory T cell Dysfunction

**DOI:** 10.1101/2022.12.02.518871

**Authors:** Tomokazu S. Sumida, Matthew R. Lincoln, Liang He, Yongjin Park, Mineto Ota, Helen A. Stillwell, Greta A. Leissa, Keishi Fujio, Alexander M. Kulminski, Charles B. Epstein, Bradley E. Bernstein, Manolis Kellis, David A. Hafler

**Affiliations:** Departments of Neurology and Immunobiology, Yale School of Medicine, New Haven, CT, USA; Broad Institute of MIT and Harvard, Cambridge, MA, USA; Computer Science and Artificial Intelligence Laboratory, MIT, Cambridge, MA, USA; Biodemography of Aging Research Unit, Social Science Research Institute, Duke University, Durham, NC, USA; Department of Allergy and Rheumatology, Graduate School of Medicine, The University of Tokyo, Tokyo, Japan; Dana-Farber Cancer Institute and Harvard Medical School, Boston, MA, USA

## Abstract

Autoimmune diseases, among the most common disorders of young adults, are mediated by genetic and environmental factors. While CD4^+^Foxp3^+^ regulatory T cells (Tregs) play a central role in preventing autoimmunity, the molecular mechanism underlying their dysfunction is unknown. Here, we performed comprehensive transcriptomic and epigenomic profiling of Tregs in the autoimmune disease multiple sclerosis (MS) to identify central transcriptional programs regulating human autoimmunity. We discovered that upregulation of a primate-specific short *PRDM1* isoform (*PRDM1-S*) induces *SGK1* independent from evolutionally conserved long *PRDM1*, leading to destabilization of Foxp3 and Treg dysfunction. This aberrant *PRDM1-S/SGK1* axis is shared among other autoimmune diseases. Furthermore, by chromatin landscape profiling in MS Tregs we identified a *PRDM1-S* specific *cis*-regulatory element associated with enriched binding of AP-1/IRF transcription factors. Our study identifies evolutionally emerged *PRDM1-S* and epigenetic priming of AP-1/IRF as key drivers of pathogenic Treg programs leading to human autoimmune disease.

## Introduction

Human autoimmune diseases constitute a leading cause of death among young adults with an increasing incidence in recent years. Multiple sclerosis (MS) is a canonical, genetically mediated autoimmune disease induced by environmental factors where genetic perturbation of *cis*-regulatory elements in pathogenic immune cells leads to immune dysregulation and generation of autoreactive T cells and antibodies^1–3^. Among immune cells, CD4^+^ T cells play a central role in both mediating and regulating autoimmunity. As CD4^+^ T cells display a large degree of functional diversity, interrogation of total CD4^+^ T cell populations has not to date identified causal transcriptional changes^4, 5^. Thus, we hypothesized that interrogation of CD4^+^ T cell subpopulations is required to elucidate the pathophysiological characteristics of CD4^+^ T cells in autoimmune disorders, allowing identification of central transcriptional factors associated with loss of immune regulation

Human CD4^+^ Foxp3^+^ regulatory T cells (Tregs) play a central role in the maintenance of immune homeostasis and prevention of autoimmunity^6–8^. We first demonstrated that Tregs from patients with autoimmune disease exhibit a dysfunctional phenotype^9–11^ and this has been subsequently found among multiple autoimmune disorders. Recent evidence has shown that environmental factors, such as vitamin D and fatty acids affect Treg phenotype and function. In addition, high salt intake has been epidemiologically linked with autoimmune diseases^12^ and higher physiologic salt concentrations induce proinflammatory Th17 cells mediated by SGK1^13^ and modulate the stability of Tregs^14, 15^, resembling the phenotype observed in autoimmune diseases including MS^16^. These environmental factors are known to act, in part, through epigenetic changes that can be identified by examining the histone and methylation landscape of cells^17^. A recent investigation of the genetic architecture of 20 autoimmune diseases and MS showed that causal variants are enriched in regulatory elements that are active in immune cells^3^. Moreover, many of these genetic loci are common among multiple autoimmune diseases, implicating the shared immunomodulatory mechanism in human autoimmunity^18, 19^. These findings, together with increases in the incidence of autoimmune diseases over the past three decades that cannot be explained by genetic factors alone, points to both genetic and epigenetic factors as key mediators of risk for autoimmunity. While these lines of evidence strongly suggest that both genetic and epigenetic alterations might disturb Treg homeostasis, the underlying mechanism in Treg dysfunction in human autoimmune diseases has not been elucidated.

We interrogated the phenotypic and functional characteristics of human CD4^+^ T cells, focusing primarily on memory Treg (mTreg) through comprehensive transcriptomic and epigenetic profiling. We adopted both bulk and single-cell RNA-seq (scRNA-seq) for transcriptomic profiling and Assay for Transposase-Accessible Chromatin using sequencing (ATAC-seq) for probing epigenetic regulation to understand the molecular mechanisms that drive dysfunctional programs in MS Tregs. Here, we reveal that *PRDM1*, which encodes Blimp1, is upregulated in both mTreg and mTconv from patients with MS with a more significant increase in mTreg. This transcriptional signature of Tregs is shared across different autoimmune diseases, suggesting the common transcriptional signature driving dysfunctional Tregs in human autoimmunity. Specifically, an alternative short isoform of *PRDM1 (PRDM1-S)*, which codes for protein in primates but not in rodents, is primarily elevated in MS mTreg compared to the evolutionary conserved long *PRDM1* isoform (*PRDM1-L*). Gene overexpression experiments in primary human Tregs, together with bulk and single-cell RNA-seq transcriptional profiling, demonstrate a unique link between the *PRDM1-S* and *SGK1* that accounts for Treg dysfunction observed in MS. Moreover, while genome-wide chromatin accessibility in mTreg remains comparable between MS and healthy controls, the transcription factor (TF) footprints and motifs within accessible chromatin regions are different with significantly enriched binding of AP-1 and IRF family TFs observed in MS mTreg, suggesting rewiring of regulatory circuits. Finally, CRISPR activation (CRISPRa)-based exploration of *cis*-regulatory elements for the *PRDM1-S* identified a *cis*-regulatory element harboring AP-1 and IRF4 composite motif. These findings suggest a possible novel regulatory program by which Tregs become dysfunctional in humans that is shared across multiple autoimmune diseases. Our multimodal datasets of human CD4^+^ T cells provide a rich resource for understanding the loss of immune regulation in autoimmune diseases and suggest that the primate specific short *PRDM1* isoform is a critical, targetable transcriptional regulator in human autoimmunity.

## Results

### Deep transcriptomic analysis of memory Treg and Tconv highlight PRDM1 upregulation in MS (Figure 1)

We sought to identify CD4^+^ T cell transcriptional differences between patients with MS and healthy controls. As previous studies had not identified significant differences in bulk transcriptional profiles of whole CD4^+^ T cells between MS subjects and healthy controls ^4, 5^, we divided CD4^+^ T cells into four major subpopulations based on two categories; Tconv vs. Treg, and naïve vs. memory (Figure S1A), where each population is hypothesized to be involved in MS pathogenesis. For example, mTconv contains pathogenic CD4^+^ T cells in patients with MS, such as myelin-reactive T cells, which display the signatures of Th1 and/or Th17 cells^20–23^. Moreover, mTregs with reduced suppressive function are implicated in MS pathophysiology^24, 25^. To provide a comprehensive transcriptomic catalogue of memory CD4^+^ T cell subpopulations in patients with relapsing-remitting MS (RRMS), we performed RNA-seq on *ex vivo* mTreg and mTconv isolated from the peripheral blood of untreated RRMS patients free of steroid treatment and healthy control subjects as a discovery cohort (Figure 1A, S1A). The control subjects were matched by age, sex, and ethnicity (clinical characteristics are described in Supplementary Table 1). Among a total of 90 RNA-seq samples (48 mTconv and 42 mTreg samples) in the discovery cohort (HC; n = 21, MS; n = 30), 84 samples (HC; n = 20, MS; n = 26) passed quality control (Method Details). We identified 21 and 243 differentially expressed genes (DEGs, defined as |log2FC| >0.6, FDR<0.1) between MS and healthy subjects for mTreg and mTconv, respectively (Figure 1B). Several DEGs were up- or down-regulated in the same direction in both mTreg and mTconv: *PRDM1*, *BCL3*, and *PIM3* were upregulated genes, and *ID3*, *TOB2*, and *LBH* were downregulated genes in MS (Figure 1C, S1B). *PRDM1* was identified as one of the top genes significantly upregulated in MS in both mTreg and mTconv (Figure S1C). Reduced expression of *ID3* in both cell types in MS reflects the negative regulation of *ID3* by *PRDM1* consistent with the known function of *ID3* in maintaining Foxp3 expression in Treg^26^ (Figure 1D). We confirmed that the frequency of Foxp3^+^ Tregs within CD4^+^ T cells were not changed in patients with MS (Figure S1D), consistent with previous studies that quantitative loss of Foxp3^+^ Tregs is unlikely to be the primary feature of disturbed peripheral T cell tolerance in patients with MS or other autoimmune diseases.

**Figure 1.**
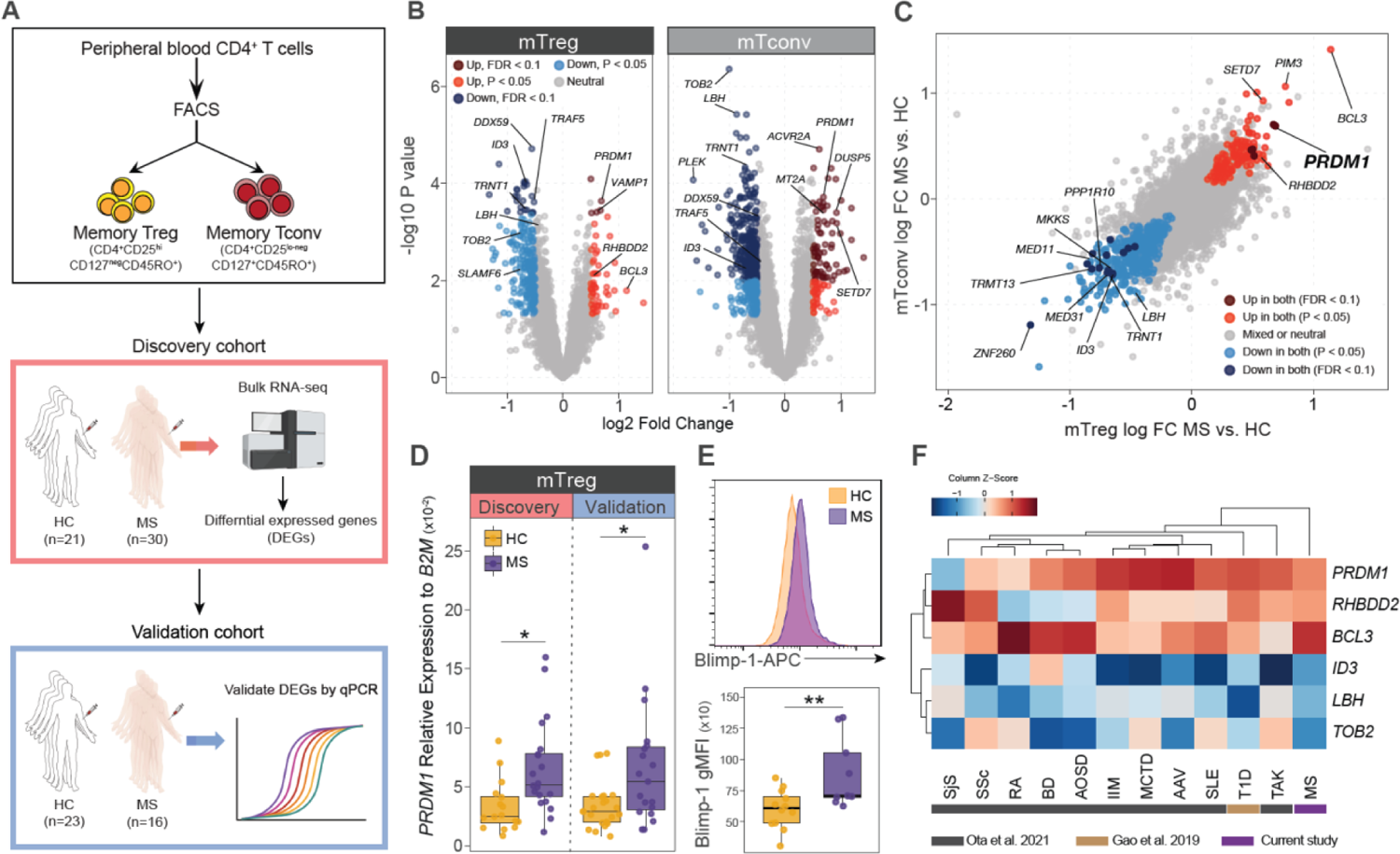
Deep transcriptomic analysis of memory Treg and Tconv highlight PRDM1 upregulation in MS. **(A)** Schematic of study design. Memory Tconv (mTconv) or memory Tregs (mTreg) were FACS sorted from peripheral blood CD4***^+^*** T cells from patients with MS and healthy control subjects (HC). Bulk RNA-seq was performed on the discovery cohort and DEGs were identified (HC: n=20, MS: n=26). The selected DEGs were validated by an independent validation cohort (HC: n=23, MS: n=16). **(B)** Volcano plots showing DEGs for mTreg and mTconv between MS and HC. **(C)** Overlapped DEGs between mTreg and mTconv. **(D)** qPCR validation for *PRDM1* expression in both discovery and validation cohorts. **(E)** Protein validation for Blimp1 expression using flow cytometry (HC; n=12, MS; n=9). **(F)** Heatmap depicting expression patterns of selected six genes in mTreg/Fr2 eTreg across 12 autoimmune diseases (data were extracted from M. Ota *et al.* and P. Gao *et al.*). P*<0.05, P**<0.01; Statistical significance computed by unpaired t-test **(D, E)**.

We validated several of the top DEGs (*PRDM1, BCL3, RHBDD2, TOB2, LBH*) in mTreg by using qPCR with an independent cohort of patients with MS (n = 16) and healthy controls (n = 23) (Figure 1A and E, Figure S1E, patient characteristics are described in Supplementary Table 1). We further confirmed the up regulation of *PRDM1* in MS mTreg and mTconv at the protein level by intracellular staining of Blimp1 using flow cytometry (Figure 1E). We then asked whether this Treg transcriptional signature observed in MS is shared across different autoimmune diseases using ImmuNexUT data^27^ where transcriptomic profiles of multiple peripheral immune cells, including mTreg and mTconv, were explored at population scale across multiple autoimmune diseases. Notably, the transcriptional signature that we observed in MS Treg was also observed in Treg among most of the autoimmune diseases we analyzed (total 12 diseases: 10 diseases from ImmuNexUT, T1D from P. Gao *et al.*^28^, and MS from this study) (Figure 1F, S1F). Inverse regulation of *PRDM1* and *ID3* observed in MS was highlighted in Tregs from patients with systemic lupus erythematosus (SLE), Idiopathic inflammatory myopathy (IIM), and ANCA-associated vasculitis (AAV) (Figure S1G). Our bulk RNA-seq transcriptional profiling of memory Tconv and Treg highlighted *PRDM1* as a key regulatory factor in dysfunctional Tregs in autoimmune diseases.

### Single-cell dual omics analysis reveals elevated PRDM1 in Th17-like Treg in MS. (Figure 2)

To gain a deeper understanding of cellular heterogeneity and novel cell types underlying disease mechanisms, we designed and performed single-cell RNA-sequencing (scRNA-seq) to profile CD4^+^ T cells. To overcome the sparsity of scRNA-seq data^29, 30^, we used Cellular Indexing of Transcriptomes and Epitopes by sequencing (CITE-seq)^31, 32^, and profiled 44 surface protein markers (Supplementary Table 2) and mRNA expression simultaneously in total CD4^+^ T cells enriched with Treg cells. Additionally, to avoid experimental batch effects between controls and MS, we adopted hashing technology to pool cells into a single run of the 10x Genomics platform. Across five experimental batches, we obtained a comparable number of Treg and Tconv cells (Figure S2A). Demographic backgrounds (age, sex, and ethnicity) are controlled in each batch of 10x Genomics processing between controls and MS, and a total of five paired MS and control samples were included (Supplementary Table 1).

**Figure 2.**
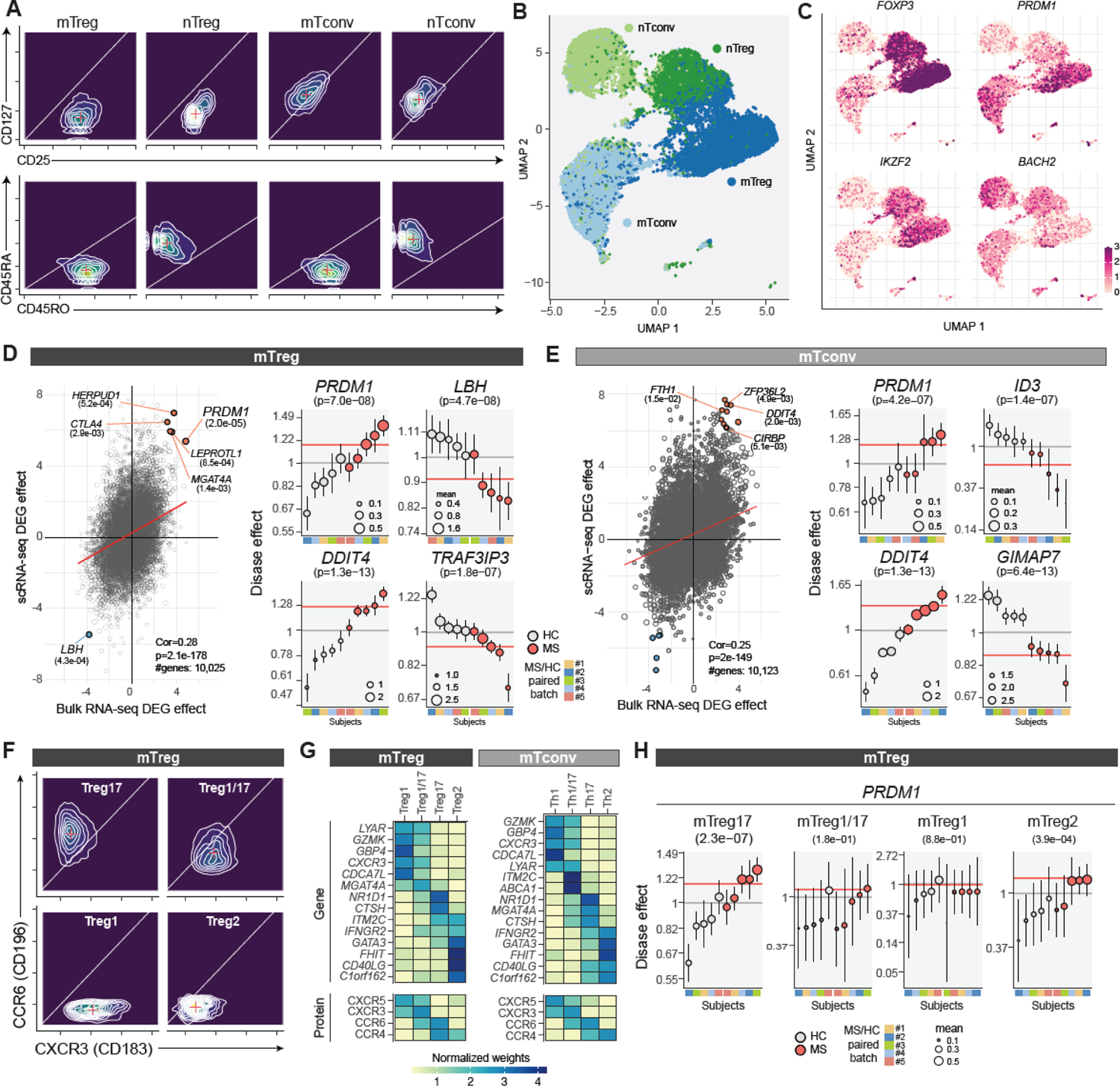
Single-cell dual omics analysis reveals elevated PRDM1 in Th17-like Treg in MS. **(A)** Surface protein guided CD4***^+^*** T cells subtype annotation. Four CD4***^+^*** T cell subtypes were distinguished by CD25, CD127, CD45RO, and CD45RA protein expression. **(B)** UMAP based on gene expression for CD4***^+^*** T cells demonstrating decent overlap with protein-based subtype annotation. **(C)** *FOXP3*, *IKZF2, PRDM1*, and *BACH2* expressions on UMAP (all cells passed QC are plotted). **(D)** Combined differential analysis for bulk- and scRNA-seq in mTreg. Representative differentially expressed genes with pseudo-bulk analysis with scRNA-seq are shown. Gene expression changes in MS relative to control with indicated genes are computed at the single-cell level (see detailed in Methods). The size of dots is scaled proportionally to the average number of mRNA reads quantified within each batch and condition; the y-axis (disease effect) shows average gene expression after adjusted by confounding factors by matching; the error bars capture one standard deviation for the disease effects in Bayesian inference. Experimental batches for paired HC and MS samples are color coded (#1-5). **(E)** Combined differential analysis for bulk- and scRNA-seq in mTconv. **(F)** Sub-cell type analysis based on CITE-seq. Surface CXCR3 (CD183) and CCR6 (CD196) protein expressions of four subtypes are shown (log10 scale). **(G)** Heatmaps showing the marker genes and proteins to define subtypes for each mTconv and mTreg. **(H)** The changes of *PRDM1* expression at subtype-level analysis in mTreg between MS and control subjects.

We established cell identities based on surface marker proteins, eliciting prior knowledge of cell-type-specific signatures. We first built a data matrix of 25,267 features (proteins and genes) and 36,983 cells, combining five batches of CITE-seq profiles. We conducted a basic quality control procedure to remove low-quality cells (Figure S2B) followed by batch normalization to control batch-specific bias (Methods). We annotated cell type identities in semi-supervised training guided by combinations of surface markers (Figure 2A), clearly demonstrating distinctive cell populations in both the protein and transcriptomic space as can be seen in the gene-expression-based clustering patterns in the UMAP visualization (Figure 2B). We confirmed that cell type annotation results were not affected by batch labels or disease groups (Figure S2C). Our experimental design also consistently enriched rare Treg populations and provided sufficient cells to study the variation within Tregs (Figure S2D). The markers for Treg vs Tconv (*FOXP3, IKZF2*, *IL2RA/*CD25, and *IL7R*/CD127), and memory vs naive (CD45RA, CD45RO) clearly distinguished four CD4^+^ T cell subpopulations (Figure 2C, Figure S2E).

For differentially expressed gene^33^ analysis between MS and control subjects at the pseudo-bulk level^34^, we quantified subject-level gene expression profiles after adjusting contributions of putative confounding factors, which stem from unmeasured technical covariates but may create spurious associations with disease status and cell types (see Methods for details)^35^. We found that single-cell-based analysis identified 90 up-regulated genes, and 16 down-regulated genes in MS mTreg (Figure S2F). We compared the DEG effect size calculated in the scRNA-seq data (y-axis) with the bulk DEG results (x-axis) (Figure 2D and E), which confirmed that a substantial number of DEGs found in the bulk RNA-seq analysis are replicated in the matched subpopulations (mTreg and mTconv) with a statistically significant correlation in both mTreg and mTconv (mTreg; r = 0.28, p=2^-178^, mTconv; r = 0.25, p=2^-149^). For example, upregulation of *PRDM1* and downregulation of *LBH* were demonstrated in MS mTreg (Figure 2D). *PRDM1* was also upregulated in mTconv, which further validated the importance of *PRDM1* in the MS T cell signature (Figure 2D and E, Figure S2G). *DDIT4*, which suppresses mTOR function, was upregulated in mTreg as well as mTconv and downregulation of *TRAF3IP3* in MS mTreg, supporting the dysfunctional Treg property and skewed Th17 signature in MS^36–39^. Of note, expression of CD45RO and *FOXP3* was not significantly altered between MS and control subjects (Figure S2G), indicating our differential analysis based on dual omics single-cell analysis was not biased by skewed memory differentiation or significant loss of *FOXP3* gene expression in MS mTreg.

To elucidate the plasticity of the memory CD4^+^ T cell population, we further determined the T helper cell subtypes (Th1, Th2, Th17, Th1/17) in mTreg and mTconv by using key marker expression with CITE-seq (Figure 2F, G, Figure S3A-E, see also Methods). We then analyzed the gene expression changes between MS and controls at each mTreg subtype and found that *PRDM1* was significantly upregulated in Th17-like mTreg (mTreg17) (Figure 2H). Upregulation of CD226 and *CD6* in mTreg17 and CD278 (ICOS) and *JUNB* in Th17 indicates the skewed pathogenic Th17-like signature in MS memory T cells (Figure S3F)^40–43^. Given the causative role of Th17 in MS pathophysiology, our results at single-cell resolution further dissected the features of MS mTreg and highlighted the potent role of *PRDM1* in skewing mTreg and mTconv toward the pathogenic Th17-like signature in the context of MS.

### Elevated alternative short PRDM1 isoform in MS mTreg. (Figure 3)

Our unbiased transcriptomic profiling of MS mTreg using bulk and single-cell RNA-seq identified *PRDM1* as a top candidate transcription factor accounting for dysfunctional Treg properties in MS. *PRDM1* encodes the Blimp1 protein which functions as a zinc finger transcriptional repressor initially identified as a protein that binds to the promoter of *IFNB1* and suppresses its expression^44^. The development and function of a variety of immune cells are also under the control of Blimp1. CD4^+^ T cell-specific *PRDM1* deletion exaggerates the proinflammatory reaction in multiple murine models of autoimmune diseases, including EAE^45, 46^; in contrast, previous studies demonstrated Blimp1 as an essential factor driving inflammatory programs in Th17 differentiation^47^. This contradictory evidence suggests the context-dependent roles of *PRDM1* in CD4^+^ T cells in mice. Recently, the role of *PRDM1* in Treg cells was studied by using Treg-specific *PRDM1* knockout mice in the context of autoimmunity. Loss of their suppressive capacity in *PRDM1*-deficient Treg indicates that *PRDM1* plays a critical role in maintaining Treg homeostasis in tissue and positively regulates its suppressive function^48–50^. Thus, our result showing upregulation of *PRDM1* in MS mTreg fundamentally contradicted observations in mice.

**Figure 3.**
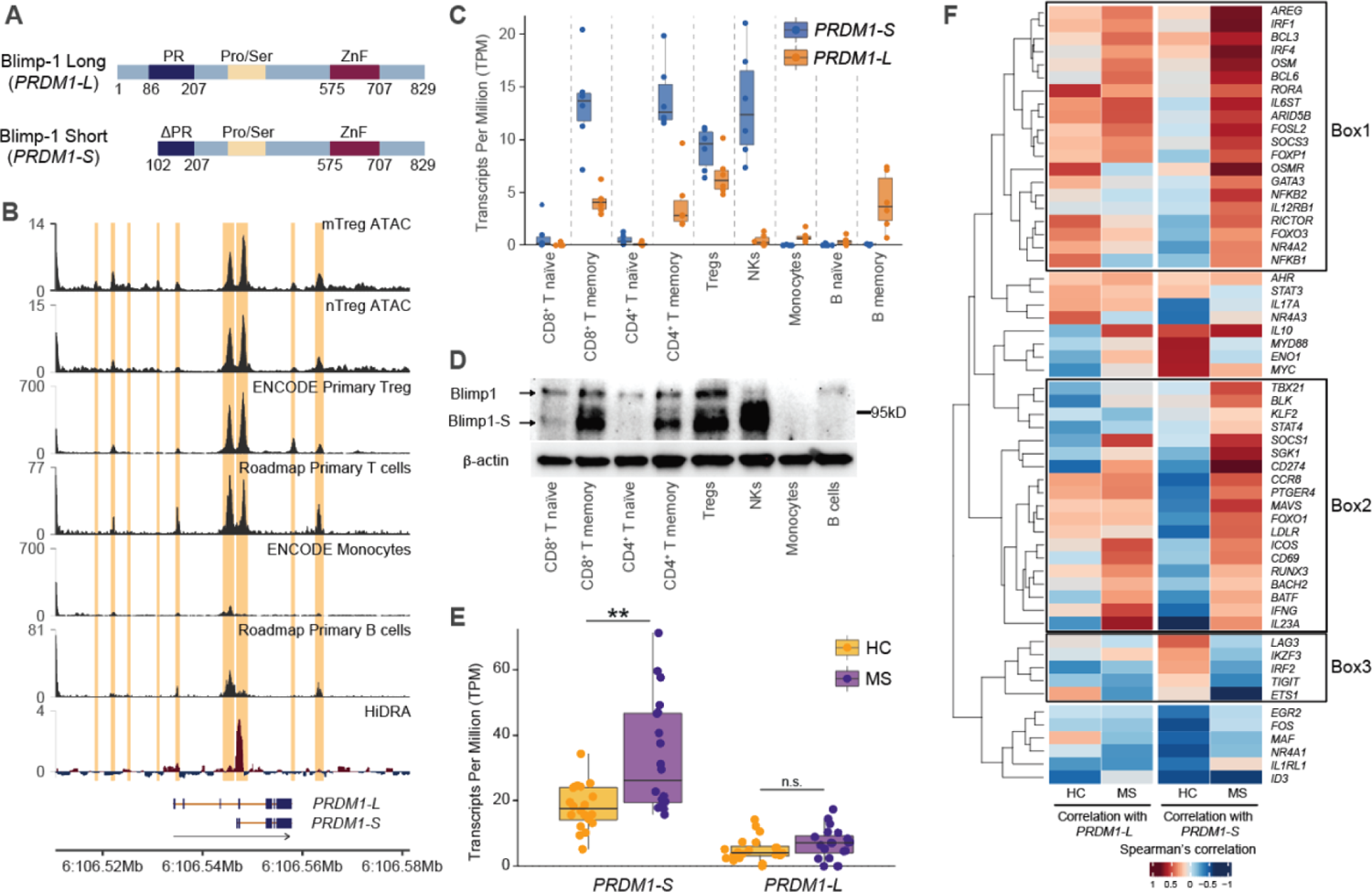
Elevated alternative short PRDM1 isoform in MS mTreg. **(A)** Schematic of *PRDM1* short and long isoforms. PR; PR domain, Pro/Ser; Proline/serine rich region, ZnF; five C2H2 zink fingers. **(B)** ATAC-seq (mTreg and nTreg), DHS (ENCODE primary Treg, Roadmap primary T cells, ENCODE monocytes, Roadmap Primary B cells), and HiDRA peaks at *PRDM1* locus. **(C)** *PRDM1-S* and *PRDM1-L* isoform expression across 9 different immune cell types in peripheral blood assessed by bulk RNA-seq (n=6). **(D)** Western blot analysis of Blimp1 expression from 8 different immune cell types in peripheral blood. Conventional Blimp1 and alternative Blimp1-S are distinguished by different size. **(E)** *PRDM1-S* and *PRDM1-L* gene expression assessed by bulk RNA-seq in mTreg between MS and HC. P**<0.01; Statistical significance computed by one way ANOVA with Dunn’s multiple comparisons tests. **(F)** Heatmaps depicting the Spearman’s correlation for curated immune related genes with *PRDM1-S* or *PRDM1-L* in HC or MS.

In humans, it has been appreciated that *PRDM1* has two major isoforms: the original full-length isoform and another short isoform generated by alternative promoter usage^51^. The short *PRDM1* isoform (*PRDM1-S;* encoding a short form of Blimp1, Blimp1-S) arose during dry-nosed primate evolution, and thus is not coded in the mouse genome. Blimp1-S lacks the N-terminal region of Blimp1, which results in missing a part of the PR domain that is important in mediating interaction with co-factors of Blimp1 (Figure 3A). Thus, Blimp1-S is implicated as a dominant negative form against the full-length Blimp1^51, 52^. Chromatin accessibility analysis at the *PRDM1* locus showed that the *PRDM1-S* promoter region is significantly accessible in human Treg and our previous HiDRA analysis^53^ revealed stronger activity at the promoter of *PRDM1-S* than that of full-length *PRDM1* isoform (*PRDM1-L*), which is consistent with DNase I hypersensitive site (DHS) data of human primary total Treg^54^ and T cells^55^ (Figure 3B). In addition, bulk RNA-seq with nine different peripheral blood human immune cells confirmed that both short and long *PRDM1* isoforms were expressed in a cell-type-dependent manner. More specifically, monocytes and B cells mainly express *PRDM1-L*. In contrast, NK cells dominantly express *PRDM1-S*, and CD4^+^ T cells (including Treg) and CD8^+^ T cells express more *PRDM1-S* than *PRDM1-L*, especially in the memory population, suggesting cell type-specific roles for each isoform (Figure 3C, S4A). This cell-type specific expression patterns of *PRDM1-S* and *-L* were further validated at protein level by western blot (Figure 3D, S4B). There are three bands detected between 80-110 kD consistent with previous studies^56^, though the band sizes are slightly higher in our data. To confirm the band size of Blimp1-S and Blimp1 protein, we overexpressed the open reading frame of Blimp1-S or Blimp1 in 293T cells by lentiviral transduction and found that lower two bands are derived from Blimp1-S while the higher molecular weight band is from Blimp1 (Figure S4B). We then analyzed our mTreg bulk RNA-seq at transcript level and found that *PRDM1-S* is significantly increased in MS mTregs compared to healthy controls, though there is a moderate difference for *PRDM1-L* expression between MS and control subjects (Figure 3E). This alteration was further validated by qPCR in both discovery and validation cohorts (Figure S4C). Notably, this upregulation of *PRDM1-S* expression was also observed in SLE Tregs from ImmuNexUT data^27^ (Figure S4D). These data suggest that *PRDM1-S* is a key regulator of Treg function in autoimmune diseases. These lines of data prompted us to hypothesize that the aberrant induction of *PRDM1-S* may confer dysfunctional properties to MS Tregs. Given that Blimp1-S can serve as a dominant negative isoform against conventional Blimp1, we first examined whether the ratio of *PRDM1-S* and *PRDM1-L* is altered between MS vs HC. However, there was no significant difference observed in this ratio with either bulk RNA-seq or qPCR data, suggesting that the balance between *PRDM1-S* and *PRDM1-L* was not significantly disrupted in MS Tregs (Figure S4E). Although *PRDM1-S* level was significantly upregulated in MS Tregs, *PRDM1-L* levels were not changed or slightly increased in MS Treg (Figure 3E, S4C), thus *PRDM1-S* mediated effects in MS Tregs could be independent from its dominant negative function against *PRDM1-L*.

In order to identify genes that are differentially correlated between *PRDM1-S* and *PRDM1-L*, we further analyzed the co-expression pattern of immune-related genes with *PRDM1-S* and *PRDM1-L* in mTreg from HC and MS (Figure 3F); genes associated with effector and tissue resident Treg signature (i.e. *BATF, CCR8, ICOS, CD69, IL1RL1, AREG, IRF4*) were co-expressed with both *PRDM1-L* and *PRDM1-S,* particularly in MS Tregs, which reflects the skewing feature of MS mTreg towards effector/tissue resident properties. Differentially expressed genets in MS mTregs, such as *BCL3,* were positively correlated with both *PRDM1* isoforms in MS but negatively correlated in healthy controls. Of note, these positive correlations in MS are stronger for *PRDM1-S* compared to *PRDM1-L*. This type of correlation pattern (positive correlation for MS but negative for control Tregs and higher correlation with *PRDM1-S* than *PRDM1-L*: Box 1 and 2 in Figure 3F) was observed for the following genes; *IRF1*, which acts as a negative regulator for Foxp3 expression^57^; *BATF* and *FOSL2*, which belong to AP-1 family, with crucial functions for Treg differentiation and maintenance^58, 59^; *SGK1*, which is known to disrupt Treg homeostasis^14, 15^ and play pathogenic roles in EAE^60^. We also noted that suppressive molecules enhancing Treg function (i.e. *IKZF4, TIGIT, LAG3, ID3*) are negatively correlated with *PRDM1-S* in MS but not in healthy controls (Box 3 in Figure 3F), supporting the association between *PRDM1-S* and dysfunctional Treg properties in MS. Taken together, we identified *PRDM1-S* as a significantly upregulated alternative isoform that partially accounts for the dysfunctional gene signature of MS mTreg.

### Short PRDM1 induces SGK1 and Treg dysfunction. (Figure 4)

Next, we hypothesized that *PRDM1-S* has a unique feature independent of interfering from *PRDM1-L* function. To test this hypothesis, we transduced *PRDM1-S* or *PRDM1-L* into primary human Tregs by using a lentivirus-based overexpression (OE) system. Isolated human primary Tregs were infected with lentivirus encoding *PRDM1-S* or *PRDM1-L* with GFP reporter and GFP positive transduced cells were sorted by FACS after four days of culture. Overexpression for each transcript was confirmed by qPCR (Figure S5A). Since only three amino acids are unique to *PRDM1-S* (Blimp1-S) coding sequence compared to *PRDM1-*L (Blimp1), overexpression of Blimp1-S open reading frame cannot be detected by qPCR, because qPCR primers are targeted to unique sequence of 5’ UTR region of *PRDM1-S*. Instead, we confirmed upregulation of total *PRDM1* but not *PRDM1-L* with Blimp1-S OE by qPCR (Figure S5A). Protein level induction of Blimp1-S with this system was confirmed by western blot (Figure S4B). Bulk RNA-seq was performed on sorted GFP positive cells and highlighted 100 genes exhibiting nominal evidence of differential expression (|log2FC| > 0.6, P value < 0.05) between *PRDM1-S* overexpression and GFP control (Figure 4A). We found that *SGK1* was one of the major upregulated genes induced by *PRDM1-S* but not by *PRDM1-L* overexpression (Figure 4A, S5B). We validated this observation by qPCR in human primary Treg cells and Jurkat T cells (Figure 4B). Moreover, *SGK1* was upregulated in our scRNA-seq analyses, particularly in the Th17-like Treg subset together with *PRDM1*, consistent with the role of SGK1 skewing pathogenic Th17 like signature in Tregs^60^ (Figure 4C, S5C). This *PRDM1-SGK1* axis was a common feature among other autoimmune diseases in a published dataset^27^, where both *PRDM1* and *SGK1* were significantly increased in Tregs from patients with SLE and ANCA-associated vasculitis (Figure S1E, S5D). Finally, we examined the impact of *PRDM1-S* on Treg function by *in vitro* Treg suppression assay and found that Tregs with *PRDM1-S* overexpression (OE) exhibited lower suppressive function than control GFP OE Treg, strongly indicating that aberrant *PRDM1-S* expression causes Treg dysfunction (Figure 4D). *PRDM1-S* OE decreased the level of both full-length Foxp3 and the exon 2-containing suppressive isoform of Foxp3 protein, further confirming the impaired suppressive function on Tregs with *PRDM1-S* OE (Figure 4E, S5E). These data support the unique role of *PRDM1-S* as a positive regulator for *SGK1* in human Treg cells, especially in the subset of potentially pathogenic Th17-like Treg, where *SGK1* plays a proinflammatory role under high sodium conditions and is responsible for pathogenic features in a murine MS model via suppressing Foxp3 expression and stablization^14, 15^. Thus, these data suggest that the *PRDM1-S/SGK1* axis underlies Treg dysfunction in MS.

**Figure 4.**
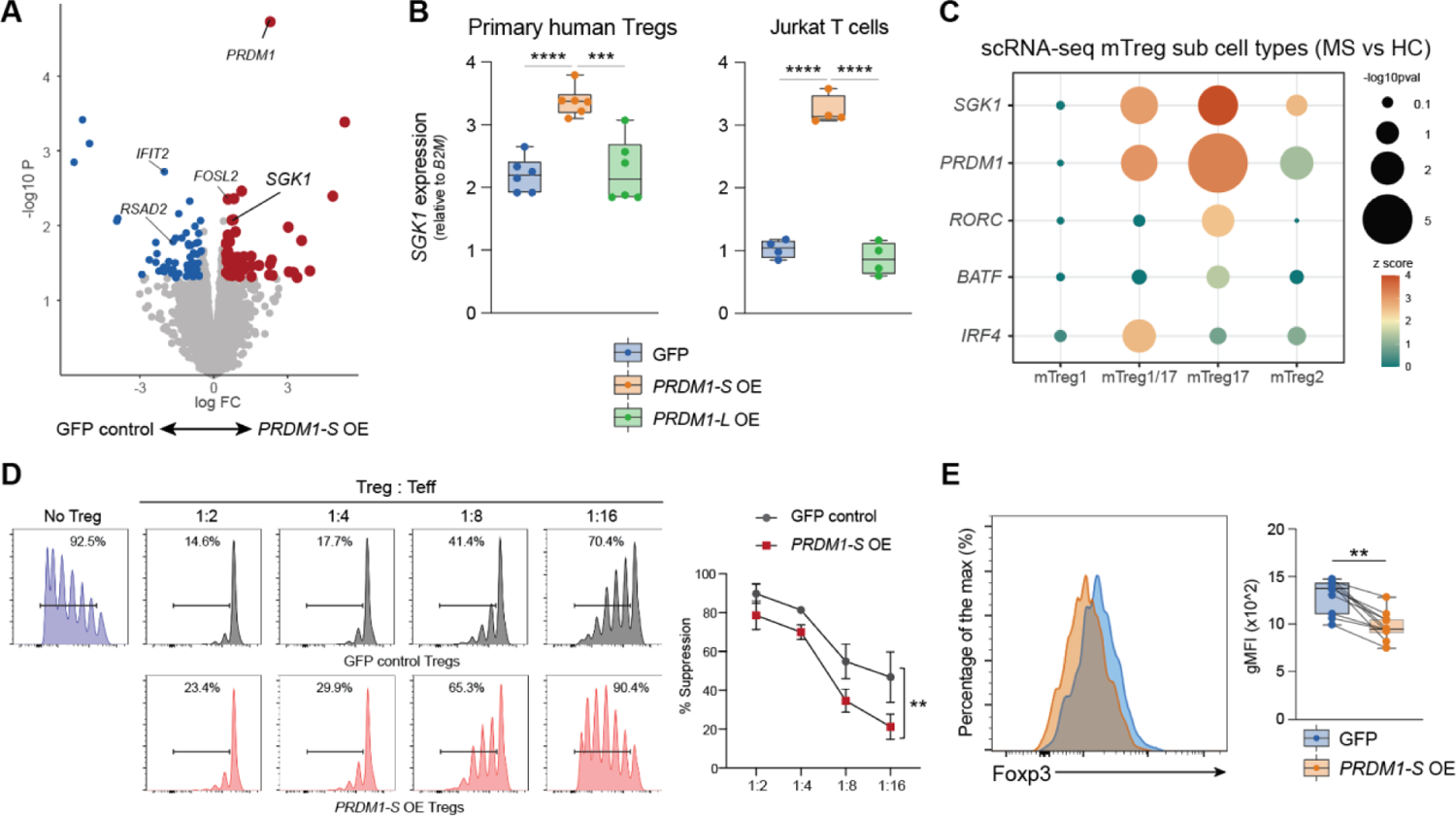
Short PRDM1 induces SGK1 and Treg dysfunction. **(A)** Volcano plot showing statistical significance and fold change for genes differentially expressed by *PRDM1-S* overexpression in primary mTreg. **(B)** *SGK1* expression was assessed by qPCR for overexpression of *PRDM1-S* and *PRDM1-L* with primary human mTregs (left) and Jurkat T cells (right). P***<0.001, P****<0.0001; Statistical significance computed by one way ANOVA with Dunn’s multiple comparisons tests. **(C)** Pseudo-bulk analysis of *SGK1, PRDM1, RORC, IRF4 and BATF* expression in scRNA-seq at each of four major mTreg subtypes. **(D)** *In vitro* Treg suppression assay with human primary Tregs. T effector cell proliferation was assessed after 5 days of co-culture with Tregs transduced with GFP control vector vs *PRDM1-S* OE vector. P**<0.01; Statistical significance computed by two-way repeated measures ANOVA. **(E)** Flow cytometry analysis for Foxp3 in primary Treg cells by overexpression of *PRDM1-S* compared to GFP control (n=10). P**<0.01; Statistical significance computed by paired t test.

### Comprehensive analysis for chromatin accessibility reveals AP-1 and IRF enrichment in MS mTreg. (Figure 5)

To further elucidate the regulatory mechanisms underlying the dysfunctional properties of MS mTregs, we performed Assay for Transposase Accessible Chromatin sequencing (ATAC-seq) to identify chromatin-accessible signatures and epigenetic regulatory elements. We first determined the change of mTreg chromatin accessibility between MS and healthy controls. To our surprise, there was no significant difference in genome-wide chromatin accessibility between MS and healthy controls in mTreg (FDR<0.05), suggesting that global chromatin accessibility is not per se the major factor regulating gene expression in MS mTreg. We hypothesized that differential binding of regulatory TFs may account for the changes in mTreg gene expression between MS and control subjects. To identify TFs that potentially drive the observed gene expression signature in MS mTreg, we analyzed the enrichment of TF motifs and TF footprints within accessible regions between MS and healthy controls in mTreg (Figure 5A). We observed an enrichment of AP-1 and IRFs TF motifs that are important for CD4^+^ T cell activation and differentiation in MS mTreg (Figure 5B).

**Figure 5.**
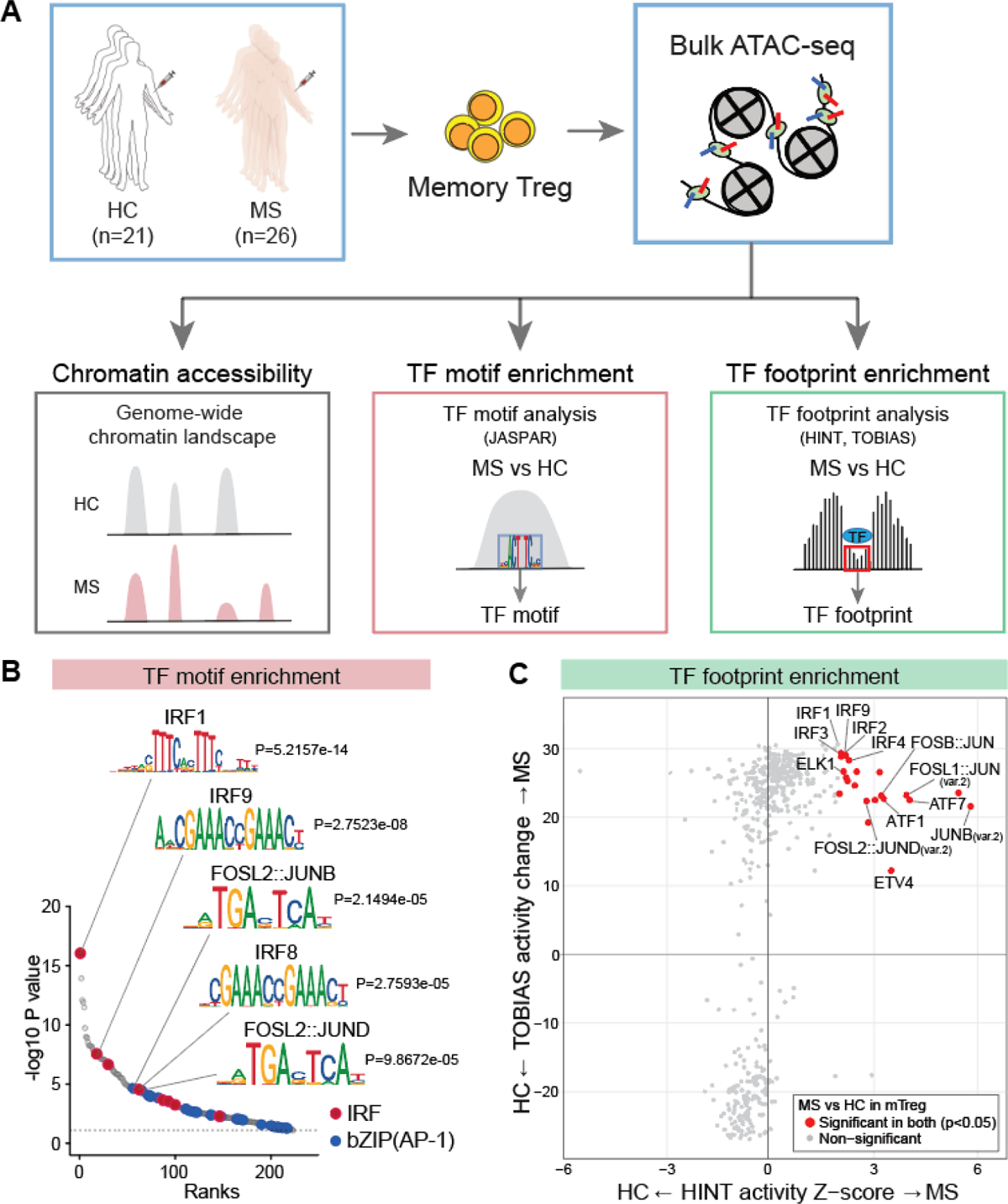
AP-1 and IRF TF bindings are enriched in MS mTreg. **(A)** Schematic of ATAC-seq experiments for mTreg from MS (n=26) and healthy control (n=21). **(B)** TF motif enrichment analysis in mTreg between MS and HC by HINT. IRF and AP-1 motifs are significantly enriched in MS mTreg. **(C)** TF footprint enrichment analysis in mTreg between MS and HC by HINT and TOBIAS. IRF and AP-1 footprints are significantly enriched in MS mTreg.

To gain deeper insight into captured accessible regulatory elements, we performed a differential footprint analysis on ATAC-seq peaks by using HINT (<u>H</u>mm-based Ide<u>N</u>tification of <u>T</u>ranscription factor footprints)^61^ and TOBIAS^62^. Consistent with our motif analysis, footprint analysis demonstrated the enrichment of AP-1 family TFs and IRFs in mTregs from MS compared to control subjects^63–66^ (Figure 5C). AP-1 transcriptional activity is negatively regulated by direct interaction with Foxp3^67^ and AP-1 is postulated to serve as a pioneer factor at Treg-specific regulatory elements where Foxp3 subsequently replaces it to establish Treg-specific enhancer architecture and DNA methylation ^68, 69^. Our observation of AP-1 enrichment in dysfunctional Tregs in MS could thus stem from the lower or impaired activity of Foxp3 in MS mTreg and reflect a more effector Tconv-like mTreg function in patients with MS^70, 71^.

IRF family TFs share a common DNA binding sequence (IRF binding element). Of interest, IRF-1 and IRF-2, but not IRF-4 nor IRF-8, are known to compete with evolutionally conserved *PRDM1-L*^72, 73^. IRF-1 plays a critical role in Treg differentiation and maintenance^57, 74^ and it was of interest that *IRF1* was more co-expressed with *PRDM1-S* as compared to *PRDM1-L* in MS mTreg (Figure 3F). In addition, the enrichment of IRF-1 TF motif and footprint in MS mTreg (Figure 5B, C) could reflect the disrupted *PRDM1-L-*mediated gene regulation in MS mTreg. These results suggest that *PRDM1-L* mediated gene regulation could be disrupted in the context of MS (Figure S6A). To test this hypothesis, we first defined the set of genes that are specifically regulated by *PRDM1-L* by performing *PRDM1-L*-specific gene knockdown in human primary Tregs. A total of 1566 differentially expressed genes (defined as |log2FC| >1, FDR<0.05; 753 upregulated and 813 downregulated genes) were identified and defined as “*PRDM1-L* signature genes” (Figure S6A and B). We performed single-cell level correlation analysis with our scRNA-seq data using a scalable negative binomial mixed model, NEBULA^75^. We then investigated putative pathogenic downstream effects of *PRDM1* expressions in single-cell analysis, examining whether the *PRDM1-L* signature genes were perturbed by *PRDM1* genes between the mTreg cell groups derived from the healthy and MS samples. We first ranked genes according to the correlation with single-cell *PRDM1* expression levels in two independent cell groups derived from the healthy and MS samples, respectively (thus, two lists of genes--one for the healthy and the MS). At a p-value threshold (the x-axes of Figure S6C), we investigated whether the number of genes in the top lists were from the *PRDM1*-L signature genes. So as not to be biased by a fixed p-value threshold, we conducted bootstrapped enrichment tests for each p-value threshold. We found that the *PRDM1*-L signature genes were significantly depleted in the top list of the genes derived from the MS mTreg group, suggesting that the interactions between *PRDM1* and the target genes were more frequently disrupted in the disease cells. We further explored the isoform level transcriptional co-regulation by using our bulk RNA-seq data. Approximately half of positively correlated *PRDM1-L* signature genes in Tregs from healthy controls lost their positive correlation in MS Tregs (Figure S6D, Supplementary Table 3). Given that *PRDM1-L* plays a crucial role in maintaining Treg function ^48–,50^, we next assessed the correlation between *PRDM1-L* and Treg signature genes^76, 77^. As observed with *PRDM1-L* signature genes, approximately half of the positively correlated Treg signature genes in healthy controls lost their positive correlation in MS Tregs (Figure S6E). Taken together, our ATAC-seq results revealed that while genome-wide chromatin accessibility was not significantly altered, differential binding of TFs (AP-1 and IRFs) to regulatory elements could serve as key upstream regulatory factors and drive dysfunctional Treg gene programs by possibly disturbing *PRDM1-L* mediated gene regulation in patients with MS.

### Identification of active enhancer for Short PRDM1 in human T cells. (Figure 6)

We sought to determine the regulatory mechanisms that induce upregulation of *PRDM1-S* as opposed to *PRDM1-L* in human mTreg. We reasoned that the enriched binding of AP-1 and IRFs exert its regulatory function through binding to *cis*-regulatory elements for *PRDM1*, especially *PRDM1-S*. To identify the *cis*-regulatory elements for *PRDM1-S*, we prioritized the accessible chromatin regions surrounding the *PRDM1* locus (+/-0.5MB window around the transcriptional start site of *PRDM1*) and examined the association between *PRDM1* expression and chromatin accessibility in our human primary T cell ATAC-seq and RNA-seq data. Twenty significant accessible regions (p<0.001) were identified as potential regulatory elements associated with *PRDM1* expression (Figure 6A). The majority of these peaks overlapped with H3K27ac ChIP-seq signals for primary human Tregs^55^, nominating them as potential enhancers. To functionally validate these accessible chromatin regions, we adopted the CRISPR activation (CRISPRa) system in Jurkat T cells that stably expresses catalytically inactive Cas9 fused to the transcriptional activator VP64 (dCas9-VP64) ^78^ (Figure 6A).

**Figure 6.**
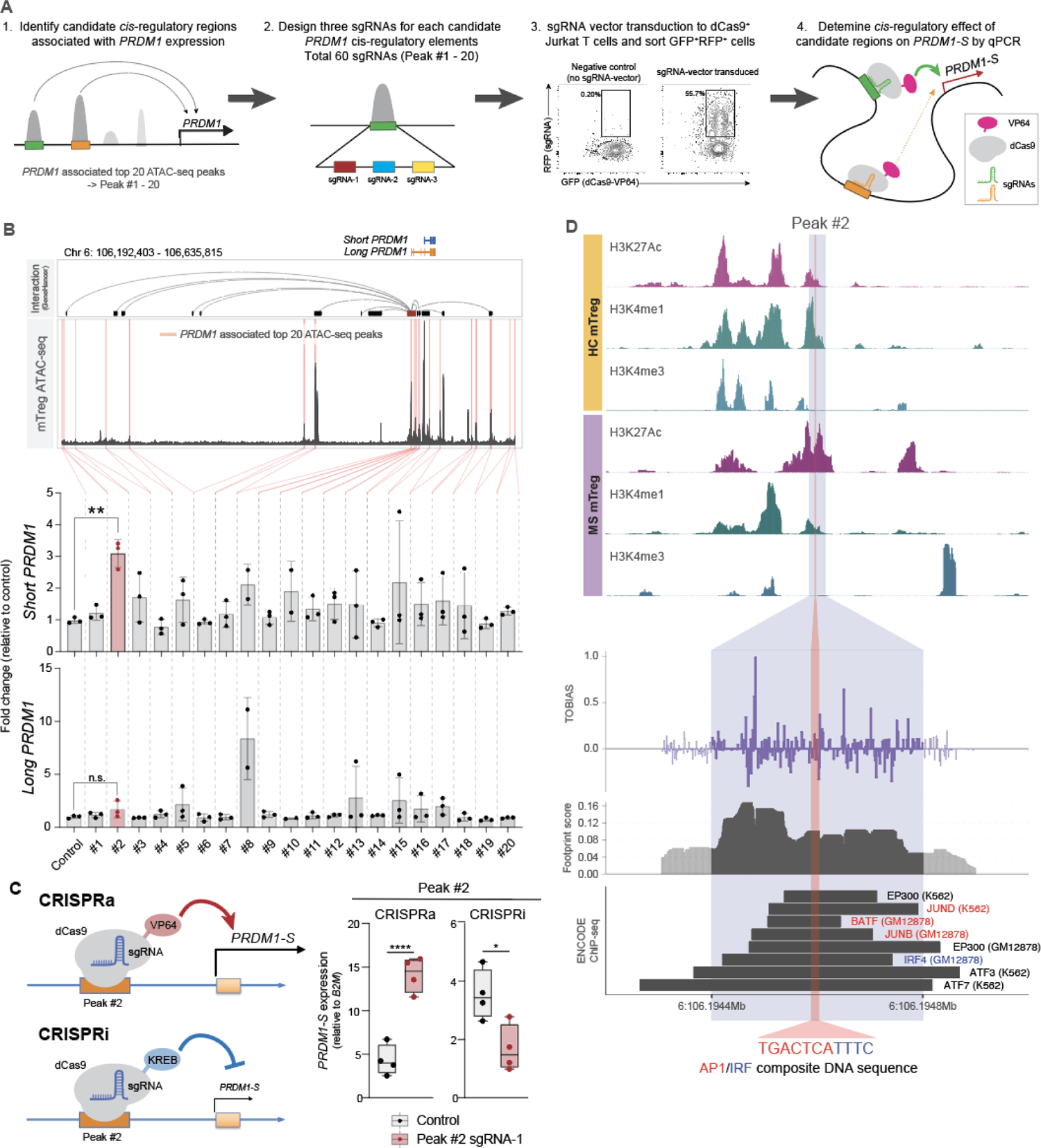
Active enhancer element for short PRDM1 with AP-1 and IRF bindings. **(A)** Schematic experimental overview. 1. Identification of candidate *cis*-regulatory elements regulating *PRDM1* expression from ATAC-seq peaks. 2. CRISPRa based examination of *PRDM1-S* specific *cis*-regulatory elements. **(B)** CRISPRa validation for top 20 *PRDM1* associated regulatory elements. Top: Top 20 accessible chromatin elements that are associated with *PRDM1* expression are highlighted in red. Potential interactions of regulatory elements with *PRDM1* gene are analyzed by GeneHancer database and shown on the top. Middle and bottom; CRISPRa-induced expression of short *PRDM1* (middle) and Long *PRDM1* (bottom) were assessed by qPCR. Detailed information for all 20 regions is shown in Supplementary Table 4. **(C)** Validation CRISPRa and CRISPRi experiment for #2 peak sgRNA-1 (n=4). **(D)** Top: H3K27ac, H3K4me1, and H3K4me3 MINT-ChIP signal on the #2 peak region in mTreg from HC and MS. Four replicates of HC and two replicates of MS are merged into one representative track respectively. Middle: Footprint analysis on #2 peak region with TOBIAS footprint score. Bottom: AP-1 and IRF ChIP-seq signals identified on #2 peak region in ENCODE data are shown. AP-1/IRF composite motif identified in #2 peak region is highlighted. P*<0.05, P**<0.01, P****<0.0001; Statistical significance computed by one way ANOVA with Dunn’s multiple comparisons tests.

First, we confirmed that *PRDM1-S* and *PRDM1-L* are independently regulated through different promoter activity by our CRISPRa method with sgRNAs targeting each promoter region, though there may be interactions between the transcription start site (TSS) of *PRDM1-L* and *PRDM1-S* (Figure S7A). Next, we designed three sgRNAs for each accessible region and generated sgRNA expressing lentiviral particles for twenty candidate regulatory elements (coded as #1 to #20) (Supplementary Table 4, Figure 6A). dCas9-VP64 expressing Jurkat T cells were infected by lentivirus encoding each sgRNA, then the double positive cells for GFP (dCas9-VP64) and RFP (sgRNA) were sorted by FACS (Figure 6A). We observed that sgRNAs targeting the #2 peak region (−339,554 bp upstream of the *PRDM1-L* TSS) mediated a unique induction of *PRDM1-S* but not *PRDM1-L* compared to control sgRNAs (Figure 6B). This #2 peak region is reported as one of the “double elite” regulatory elements for *PRDM1* in the GeneHancer dataset^79^, which reflects a higher likelihood of prediction accuracy for both enhancer and target gene (Figure 6B; top). The enhancer function of the #2 peak *cis*-regulatory element was further validated by independent experiments with not only CRISPRa but also CRISPRi (Figure 6C, S7B).

To further clarify the function of this region as a *cis*-regulatory element at *ex vivo* human primary Tregs, we sought to decode the histone modification. The main obstacle in the investigation of chromatin state by using conventional ChIP-seq technique is the limited number of *ex vivo* primary Tregs available for determining multiple histone marks from the same sample. Recent development of multiplexed, indexed T7 ChIP-seq (Mint-ChIP)^80^ allows us to identify both active and repressive epigenetic marks by using histone modification-specific antibodies with limited cell numbers. We performed Mint-ChIP on human primary Tregs in collaboration with the ENCODE project. Histone modifications (H3K27ac, H3K4me1, and H3K4me3) distinguishing active enhancers were assessed on *ex vivo* human primary Treg from MS and control subjects. The #2 peak region was marked by H3K4me1 and H3K27ac but without H3K4me3, suggesting the #2 peak region functions as an active enhancer (Figure 6D; top). Importantly, this #2 peak region overlaps with AP-1 family and IRF4 ChIP-seq peaks and contains AP-1/IRF composite motif^81^, suggesting this enhancer element regulates *PRDM1-S* via AP-1 and IRF4 binding (Figure 6D; bottom).

Given that BATF and IRF4 are known to bind in a cooperative fashion on the AP-1/IRF composite motif facilitating Th17 differentiation ^63, 82^, we examined the role of IRF4 and BATF on *PRDM1-S* expression in Tregs by performing IRF4 and BATF knockdown experiments. Surprisingly, *PRDM1-S* was upregulated by knocking down *IRF4* or *BATF* while in contrast, *PRDM1-L* was downregulated by *IRF4* KD (Figure S7C). These data indicate that *IRF4* differentially regulates *PRDM1-S* and *PRDM1-L* in human primary Tregs. Of note, *IRF4* or BATF KD did not affect *FOXP3* expression, suggesting that the loss of IRF4 and BATF in human Tregs can induce *PRDM1-S* and *SGK1* expression without significant reduction of *FOXP3* expression, further indicating that the dysfunctional *PRDM1-S/SGK1* axis observed in MS Tregs is counter regulated by the core effector Treg regulator IRF4 and BATF. These data also suggest that the *cis*-regulatory element (#2 peak region identified as upstream of *PRDM1-S*, Figure 6) serves as a negative regulatory element for *PRDM1-S* expression via IRF4/BATF binding. Indeed, CRISPRi targeting on this #2 peak region suppress *PRDM1-S* expression (Figure 6C). Thus, these results highlight IRF4/BATF as potential upstream transcription factors negatively regulating *PRDM1-S and* indicating that a newly identified *cis*-regulatory element that contains the AP-1/IRF composite motif may account for aberrant *PRDM1-S* induction in MS mTreg.

## Discussion

Disruption of peripheral CD4^+^ T cell homeostasis is a central component driving pathogenesis of autoimmune disease where autoreactive T cells lose tolerance to self-antigen by both intrinsic and extrinsic mechanisms. Treg-mediated surveillance is central for controlling activation of autoreactive CD4^+^ T cells and dysfunctional Tregs are a hallmark of MS and other autoimmune diseases ^7, 8^. In addition, recent analysis of genome-wide association studies emphasizes the substantial contribution of CD4^+^ T cells, including Tregs, as potentially causal mediators of autoimmune disease ^3^. Although several phenotypic changes have been identified in dysfunctional Tregs, the underlying molecular mechanisms leading to breakdown of Treg suppressive function in patients with autoimmune diseases are unknown. Here, by using MS as a model for studying the molecular mechanisms of human Treg dysfunction, we examined transcriptional and epigenetic alteration in human Tregs, identifying a previously unknown role of an alternative short *PRDM1* isoform in dysfunctional Treg. *SGK1* was identified as a target of short *PRDM1*, which has been reported to confer the pathogenic function of Treg in both human and mouse^14, 15, 60^. Moreover, both *PRDM1* and *SGK1* were upregulated in Tregs from different autoimmune diseases such as SLE and ANCA-associated vasculitis^27^, suggesting that the *PRDM1/SGK1* axis could serve as a common feature of Treg dysfunction in the context of human autoimmunity. Finally, exploration of epigenetic changes in MS Treg revealed an active enhancer element that induces short *PRDM1* transcription, which was validated by CRISPRa/i experiments. AP-1 family, especially BATF, and IRF4 directly bound to this regulatory element, implicating the role of these TFs in contributing to *PRDM1-S* mediated Treg dysfunction in autoimmune disease. Thus, this study provides a novel mechanistic insight of dysfunctional Tregs in the context of MS and potential therapeutic targets to reverse Treg dysfunction in autoimmune diseases.

Previous studies exploring transcriptional alterations that occur in CD4^+^ T cells in patients with MS as compared to control subjects did not identify significant differences^4, 5^. We hypothesized that it was critical to examine CD4^+^ T cell subpopulations. Thus, we further segregated CD4^+^ T cells into four subpopulations and performed transcriptional characterization using bulk RNA-seq with deeper sequencing. This allowed us to detect significant differences in gene expression between patients with MS and control subjects at both the gene and transcript level. Our bulk RNA-seq based findings in the discovery cohort were further confirmed by two different means; (1) utilization of a validation cohort with the same method as the discovery cohort, and (2) utilization of CITE-seq to determine the subpopulations and assess the difference within the third cohort, leading to highly reproducible findings. One caveat is that we were unable to quantitate *PRDM1* isoform expression at single cell resolution in our 10x genomics dataset. Full length mRNA-capturing scRNA-seq has the potential to differentiate between *PRDM1* isoforms; thus, this technique could be applied to elucidate further insight at single-cell resolution^83^. This approach will also enable characterization of further differences in T cell transcriptomics between MS and healthy subjects.

Our transcriptomic analysis of *PRDM1* isoforms identified expression patterns between *PRDM1-S* and *PRDM1-L* that are highly cell type specific. Memory T cells and NK cells expressed higher levels of *PRDM1-S* as compared to *PRDM1-L,* while monocytes/DCs, and B cells preferentially express *PRDM1-L.* Of note, *PRDM1* expression across all B cell linage (naïve, unswitched memory, switched memory, double negative B cells, and plasmablasts) is strictly limited to *PRDM1-L* over *PRDM1-S* (ImmuNexUT dataset). We also observed that the balance between *PRDM1-S* and -*L* is dynamic and changes after T cell activation and cytokine milieu *in vitro*. Thus, our findings demonstrate a tight linkage between *PRDM1* isoforms and immune pathways related to AP-1/IRFs in autoimmune diseases and highlight a fundamental role in primates for *PRDM1* isoform switching in regulating immune responses.

Epigenetic alterations in MS Tregs were assessed by using bulk ATAC-seq. Conventional characterization with differential analysis of chromatin accessibility did not reveal significant alterations of chromatin accessibility between patients with MS and healthy donors at a genome-wide level. We then used our ATAC-seq data to elucidate TF footprint enrichment and found that AP-1 and IRF family TFs are significantly enriched in MS mTreg compared to that of healthy controls. It was of interest that neither AP-1 nor IRFs were detected as differentially expressed genes by transcriptomic analysis, indicating that TF activity cannot be inferred by RNA expression. Conventional TF ChIP-seq is thought to be the best method to provide direct evidence of TF activity; however, it is technically challenging to achieve with primary human *ex vivo* mTregs due to their low frequency in circulating blood. Thus, here we employed ATAC-seq footprint analysis to overcome these issues for epigenetic profiling in human Tregs and provided novel evidence of enriched AP-1 and IRFs bindings in MS mTregs that contributing to the Treg transcriptional signature. These data agree with previous studies highlighting the important role of AP-1 in establishing epigenetic state and linking genetic susceptibility to T cell activation^84^. Of note, recent studies identified Treg specific eQTL effects with MS associated CD28 susceptible locus^85^, which was also confirmed with ImmuNexUT and DICE data (data not shown). Given that CD28 signaling is central in the activation of AP-1 function in T cells, our findings of AP-1 enrichment in MS mTregs potentially suggest that the genomic susceptibility of MS could be mediated by high activity of the AP-1 family in mTregs, resulting in increased *PRDM1-S*. Further studies focused on the functional properties of the MS-associated SNP at the CD28 locus are warranted.

A fundamental question relates to the elucidation of molecular interactions between environment triggers and gene transcription driven by allelic variation associated with disease risk that lead to autoimmune disease. We have previously shown that high Na^+^ induces SGK-1 and subsequent inactivation of Foxo1 leading to dysfunctional Tregs. This SGK1-Foxo pathway plays a role in driving pathogenic Th17 cells especially under high salt environment^22^. Other recent studies in humans have demonstrated higher Na^+^ tissue levels in a subset of patients with MS^86^. Of note, gene set enrichment analysis (GSEA) demonstrated a dysfunctional Foxo1 and Foxo3 KO Treg signature that was enriched in MS and SLE mTreg (Figure S8), consistent with our previous studies showing impaired function of Foxo1 in dysfunctional Tregs in MS^15, 87, 88^. Thus, we hypothesize that molecular interactions leading to dysfunctional Tregs in autoimmunity can be driven in part through SGK-1 mediated by high sodium concentration as an environmental factor. However, it should be pointed out that this could be just one of many environmental factors that drive autoimmune disease.

In summary, our data uncover fundamental molecular mechanisms by which Treg dysfunction is triggered in patients with MS and potentially other autoimmune diseases. Identification of the primate-specific alternative short *PRDM1* isoform induction in MS Tregs highlights the importance of studying human tissues in addition to mouse models in obtaining insight into disease pathogenesis. Enhancement of the *PRDM1/SGK1* axis in mTreg was observed in the other autoimmune diseases, suggesting shared mechanisms among dysfunctional Tregs in the context of autoimmunity. Furthermore, our findings link epigenetic priming of AP-1 and IRF TF binding in MS Tregs to short *PRDM1* induction. Finally, we believe our rich data of both transcriptome and epigenome profiles on human memory CD4^+^ T cells and Tregs will be a useful tool to explore further insights into pathogenic mechanisms of dysfunctional T cells in autoimmune diseases.

## Methods

### Study subjects and ethics statement

Peripheral blood was drawn from people with MS and healthy controls who were recruited as part of an Institutional Review Board (IRB)-approved study at Yale University, and written consent was obtained. All experiments conformed to the principles set out in the WMA Declaration of Helsinki and the Department of Health and Human Services Belmont Report.

### Human T cell isolation and culture

Peripheral blood mononuclear cells (PBMCs) were prepared from whole blood by Ficoll gradient centrifugation (Lymphoprep, STEMCELL Technologies) and used directly for total CD4+ T cell enrichment by negative magnetic selection using Easysep magnetic separation kits (STEMCELL Technologies). Cell suspension was stained with anti-CD4 (RPA-T4), anti-CD25 (clone 2A3), anti-CD45RO (UCHL1), anti-CD45RA (HI100) and anti-CD127 (hIL-7R-M21, all from BD Biosciences) for 30 minutes at 4°C. Naïve Tconv (CD4+/CD25neg/CD127+/CD45ROneg/CD45RA+), Naive Treg (CD4+/CD25hi/CD127neg/CD45ROneg/CD45RA+), Memory Tconv (CD4+/CD127+/CD45RO+/CD45RAneg), and Memory Treg (CD4+/CD25hi/CD127neg/CD45RO+/CD45RAneg) were sorted on a FACSAria (BD Biosciences). Sorted cells were plated in 96-well round-bottom plates (Corning) and cultured in RPMI 1640 medium supplemented with 5% Human serum, 2 nM L-glutamine, 5 mM HEPES, and 100 U/ml penicillin, 100 μg/ml streptomycin, 0.5 mM sodium pyruvate, 0.05 mM nonessential amino acids, and 5% human AB serum (Gemini Bio-Products). Cells were seeded (30,000-50,000/well) into wells pre-coated with anti-human CD3 (2 μg/ml, clone UCHT1, BD Biosciences) along with soluble anti-human CD28 (1 μg/ml, clone 28.2, BD Biosciences) in the presence or absence of human IL-2 (50 U/ml).

### Lentiviral transduction

#### Lentiviral production

Lentiviral plasmids encoding shRNA for gene knockdown for *PRDM1-L* or open reading frame of overexpression for *PRDM1-S* and *PRDM1-L* were obtained from Sigma-Aldrich (MISSION shRNA) and Horizon Discovery Biosciences (Precision LentiORF), respectively. dCas9-VP64-2A-GFP (Addgene 61422) and pHR-SFFV-dCas9-BFP-KRAB (addgene 46911) were used for generating Jurkat T cell lines for CRISPRa and CRISPRi, respectively. EF1a-RFP-H1-gRNA vector (CASLV502PA-R from System bioscience) was modified to introduce BsaI cut site and single sgRNAs were cloned into it by using Golden Gate Assembly kit (BsmBI-v2, New England Biolabs #E1602). All single sgRNAs used in this study are listed in Supplementary Table 5. Each plasmid was transformed into One Shot Stbl3 chemically competent cells (Invitrogen) and purified by ZymoPURE plasmid Maxiprep kit (Zymo research). Lentiviral pseudoparticles were obtained after plasmid transfection of 293T cells using TurboFectin 8.0 Transfection Reagent (Origene). The medium was replaced after 6-12 h with fresh media with 1X Viral boost (Alstem). The lentivirus containing media was harvested 72 h after transfection and concentrated 80 times using Lenti Concentrator (Origene). LV particles were then resuspended in RPMI 1640 media without serum and stored at −80°C before use. Virus titer was determined by using Jurkat T cells and Lenti-X GoStix Plus (Takara Clontech).

#### Lentiviral transduction

FACS-sorted Tregs were plated at 50,000 cells/well in round bottom 96 well plates pre-coated with anti-human CD3 (2 μg/ml, clone UCHT1, BD Biosciences) and soluble anti-human CD28 (1 μg/ml, clone 28.2, BD Biosciences), in the presence of human IL-2 (50 U/ml). After 24 h, cells were transferred into Retronectin coated 96 well plates and 25-50 μl of lenti particles were added to each well, then spinfected with high-speed centrifugation (1000 g) for 1.5 hour at 32 °C. Immediately after centrifugation, cells are placed back to the culture. On day 5, cells are harvested and GFP positive cells are sorted by FACSAria or analyzed by Fortessa. Gene knockdown effect by *PRDM1*-*L* shRNA is shown in Figure S6B.

Jurkat T cells were plated at 50,000 cells/well in round bottom 96 well plates and 25 μl of lenti particles were added to each well, and spinfected as well as above. On day 3-5, cells were scaled up to 12 well plates and followed by the second scale-up at day 7-9 into 6 well plates. Cells were stimulated with PMA and Ionomycin (50 ng/ml and 250 ng/ml respectively) for four hours and GFP^+^/RFP^+^ double positive cells were sorted directly into RNA lysis buffer by FACS Aria.

### Suppression assay

CD4^+^CD25^+^CD127^neg^ Treg cells were sorted on a FACS Aria (BD Biosciences). Treg cells were transduced with lentiviral particles containing *PRDM1-S* ORF or GFP control. GFP positive cells were sorted by FACS at day 5, and freshly sorted T effector cells were labeled with cell trace violet dye and then co-cultured with GFP^+^ Treg cells (1 x 10^4^) at different ratio with human Treg inspector beads at 2:1 bead-to-cell ratio. The proliferation of T effector cells was determined at day 4 on a BD Fortessa instrument (BD Bioscience).

### Flow cytometry analysis

Cells were stained with LIVE/DEAD Fixable Near-IR Dead Cell Stain kit (Invitrogen) and surface antibodies for 30 min at 4°C. For intracellular cytokine staining, cells were fixed with BD Cytofix ^TM^ Fixation Buffer (BD Biosciences) for 10 min at RT, then washed with PBS. Intracellular staining was performed in Foxp3 permeabilization buffer (Thermo Fisher) for 30 min at 4°C. The following antibodies were used: anti-Foxp3 (clone 150D, Biolegend, and clone PCH101, Thermo Fisher), anti-Blimp1 (clone 3H2-EB, Thermo Fisher). All antibody information is listed in Supplementary Table 6. Cells were acquired on a BD Fortessa flow cytometer and data was analyzed with FlowJo software v10 (Threestar).

### Immunoblotting

Cells were lysed with RIPA buffer containing EDTA-free cOmplete protease inhibitor cocktail (Roche) and Halt phosphatase inhibitor cocktail (Thermo Fisher Scientific). Extracted protein was quantified with a BCA kit (Thermo Scientific). 0.8-1 μg of protein extract was loaded in each lane, followed by separation by 7.5% SDS-PAGE and transfer to a nitrocellulose membrane. After 1 hour of blocking with 2.5 % bovine serum albumin (BSA) containing 1X Tris-Buffered Saline, 0.1% Tween20 Detergent (TBST), the blotted membranes were then incubated overnight with primary antibodies (anti-Blimp1 (C14A4, 1:1,000, Cell Signaling Technologies or C-7, 1:200, Santa Cruz Biotechnology), anti-β-actin (A5441, 1:10,000, Sigma-Aldrich)) in TBST with 2.5% BSA. Primary antibodies were detected by the secondary antibody horseradish peroxidase (HRP)–conjugated anti-rabbit or mouse (Cell Signaling Technology), and SuperSignal West Femto Maximum Sensitivity ECL Substrate (Pierce) was used as a chemiluminescent substrate for detecting HRP. The images were obtained with a ChemiDoc Imaging system (Bio-Rad).

### Bulk RNA-seq and ATAC-seq library preparation and sequencing

#### Bulk RNA-seq

FACS sorted cells (5,000 cells) were subjected to cDNA synthesis using SMART-Seq v4 Ultra Low Input RNA Kit for sequencing (Takara/Clontech). Barcoded libraries were generated by the Nextera XT DNA Library Preparation kit (Illumina) and sequenced with a 2×100 bp paired-end protocol on the HiSeq 4000 Sequencing System (Illumina).

#### Bulk ATAC-seq

We adopted FAST-ATAC for our FACS sorted CD4^+^ T cells (5,000 cells) (Corces et al., 2016). Cells were pelleted by centrifugation at 500 g for 7 min at 4°C, then resuspended with 50 μl of transposase mixture (25 μL of 2x TD buffer (Illumina), 2.5 μL of TDE1 (Illumina), 0.5 μL of 1% digitonin (Thermo Fisher), 22 μL of nuclease-free water). Transposition reactions were incubated at 37°C for 30 minutes in a thermal shaker with agitation at 300 RPM. Transposed DNA was purified using a MinElute Reaction Cleanup kit (QIAgen) and purified DNA was eluted in 20 μL elution buffer. Transposed fragments were amplified and purified as described previously ^89^ with modified primers ^90^. Libraries were quantified using qPCR (KAPA Library Quantification Kit) prior to sequencing. All Fast-ATAC libraries were sequenced using a 2×100 bp paired-end protocol on the HiSeq 4000 Sequencing System (Illumina).

### Mint-ChIP library preparation and sequencing

We used 800,000-1,000,000 cryopreserved sorted human primary Tregs for Mint-ChIP profiling. Cells were thawed and immediately pelleted by centrifugation at 400 g for 7 min at 4°C, then resuspended with 100 μL of ice cold PBS. Approximately 100,000 cells per antibody are lysed in detergent and chromatin is digested with micrococcal nuclease. Double stranded adapters (that contain both a promoter for transcription by T7 RNA polymerase and a demultiplexing adapter) are ligated to the chromatin. Chromatin is mixed overnight with antibodies recognizing histone modifications, and immune complexes are captured using protein A / protein G magnetic bead mixtures. In addition, some chromatin is set aside overnight to enable preparation of an antibody free, input control library. Immobilized immune complexes are washed, and immunoprecipitated DNA is eluted using proteinase K. Recovered DNA is purified with SPRI beads and subject to T7 RNA synthesis, thus creating an RNA copy of the immunoprecipitated DNA. RNA is copied back to cDNA using a random primer containing a 5’ extension, enabling subsequent PCR amplification with Illumina indexed sequencing primers. PCR products are purified and mixed together to enable multiplex Illumina sequencing. DNA is sequenced using a paired end protocol; in Mint-ChIP3, the first 8 bases of Illumina Read2 serve as an inline barcode enabling demultiplexing of the chromatin using the ligated barcoded adapter. The detailed protocol and primer/adaptor sequences are described in <u>dx.doi.org/10.17504/protocols.io.wbefaje</u>.

### Mint-ChIP data processing

We processed the Mint-ChIP FASTQ files of each sample using the ENCODE3 ChIP-seq pipeline provided by Anshul Kundaje (https://github.com/ENCODE-DCC/chip-seq-pipeline2) with the following parameters “chip.pipeline_type=histone, chip.aligner=bowtie2, chip.true_rep_only=true, chip.paired_end=true, chip.ctl_paired_end=true, chip.always_use_pooled_ctl=false” specified in its json file. We used the default value for all the other parameters. Briefly, the pipeline first mapped the reads to the hg19 human reference genome using bowtie2. The aligned reads were filtered and duplicated reads were removed. The peak calling was then performed using MACS2 with a control sample for each individual. Sample quality was assessed with a cross-correlation plot.

### Bulk RNA-seq analysis

After sequencing, adapter sequences and poor-quality bases (quality score < 3) were trimmed with Trimmomatic. Remaining bases were trimmed if their average quality score in a 4 bp sliding window fell below 5. FastQC (https://www.bioinformatics.babraham.ac.uk/projects/fastqc/) was used to obtain quality control metrics before and after trimming. Remaining reads were aligned to the GRCh38 human genome assembly with STAR 2.5.2 ^91^. We used Picard (https://github.com/broadinstitute/picard) to remove optical duplicates and to compile alignment summary statistics and RNA-seq summary statistics. After alignment, reads were quantitated to gene level with RSEM ^92^ using the GENCODE annotation ^93^.

We conducted our initial quality control assessment on the entire dataset, including memory and naïve Tconv and Treg cells obtained from MS patients and healthy controls. A subset of these were stimulated with IL-2 as described above, with the remainder collected in the *ex vivo* state. We used principal component analysis to identify potential sample swaps. We considered genes that were quantitated >1 count per million (cpm) in ≥ 15 samples, normalizing expression values by the trimmed mean of M-values as implemented in edgeR ^94^. We used limma ^95^ with the voom transformation ^96^ to identify differentially expressed genes (DEGs) within mTconv and mTreg cell populations separately. We used RUV-seq ^97^ to account for batch effect and other sources of systematic variation; we included RUV parameters along with a sex covariate in our final model. DEG list for *ex vivo* mTreg and mTconv between MS vs HC are shown in Supplementary Table 7.

### Co-expression analysis in bulk RNA-seq and scRNA-seq

The co-expression between two genes was computed as the Spearman correlation coefficient of normalized gene expression. The normalized gene expression was calculated by dividing the raw count by the library size of the sample and its scaling factor obtained from the TMM normalization. The co-expression analysis of *PRDM1* in the scRNA-seq data was conducted for genes that had the average count per cell >0.005 in the mTreg and mTconv. The co-expression was measured by the log(fold-change) between *PRDM1* and the gene of interest in a negative binomial mixed model with the subjects as random effects implemented in NEBULA ^75^. We used normalized expression of *PRDM1* (the raw count divided by the library size of the cell) as the explanatory variable and included in the model the proportion of reads from ribosomal protein genes and mitochondrial genes as covariates. To assess the differential co-expression, the model was fitted for 4896 and 4676 cells from the five MS patients and five healthy controls separately.

### Bulk ATAC-seq analysis

The official pipeline of the Encyclopedia of DNA Elements ^54^ consortium (https://github.com/kundajelab/atac_dnase_pipelines) {*kundajelab/atac_dnase_pipelines: 0.3.0*} was adopted to preprocess the ATAC-seq raw data. The preprocessing started with the paired-end ATAC-seq fastq files of each subject. More specifically, reads were trimmed for adapters using cutadapt ^98^ and mapped to the human reference genome (hg19) using bowtie2 ^99^. The output raw bam files were filtered, deduped, and converted to single-ended Tn5-shifted tagalign files, which were then used as the input for peak calling. The deduped bam files were used in the downstream motif and footprint analyses described in a later section.

We first called sample-specific narrow peaks (FDR<0.01) using MACS2 ^100^ from the tagalign file of each of the samples, separately, with the command “macs2 callpeak -f BED -g hs -q 0.01 --nomodel --shift -75 --extsize 150 -B --SPMR --keep-dup all --call-summits”. We calculated the fraction of reads in called peak regions (FRiP score) for each sample. We then called group-specific narrow peaks (FDR<0.01) by providing MACS2 with the tagalign files of all high-quality samples (FRiP>0.1) within each of the eight groups (healthy/MS mTconv, mTreg, naive Treg (nTreg), and naive Tconv (nTconv)) using the command “macs2 callpeak -t <TAGALIGN files of the high-quality samples within the group> -f BED -g hs -q 0.01 --nomodel --shift -75 --extsize 150 -B --SPMR -- keep-dup all --call-summits”. To obtain a unified set of peak regions across all groups, we merged the group-specific peak regions that were overlapping or had maximum distance <100bp using BEDTools ^101^ with the command “bedtools merge -i -d 100”.

Given the unified set of peak regions, the number of reads overlapping a peak was called for each subject using BEDTools. This raw count matrix was used for the downstream analyses. In the analysis of differential accessibility, we first filtered out peaks with low counts and normalized the counts of each subject using TMM ^94^. Then the *voom* function in the limma package ^95^ was used to perform the differential analysis between the cases and controls within each cell type with FRiP, sex and ethnicity as covariates.

### Correlation analysis between RNA-seq and ATAC-seq data

The analysis of the correlation between the accessibility of the adjacent open chromatin regions and the *PRDM1* expression was performed using a linear regression model in which the normalized *PRDM1* expression (raw count divided by the total library size and the scaling factor) was the dependent variable and the adjusted ATAC-seq peak height was the explanatory variable. The adjusted ATAC-seq peak height was the residual obtained by fitting a weighted linear model using limma-voom for the peak with FRiP as the covariate. We included 106 samples from all the cell populations and both groups and these samples had both RNA-seq and ATAC-seq measurements. We interrogated 72 ATAC-seq peaks for which *PRDM1* was annotated as the nearest gene by Homer. These peaks span from ∼350k bp upstream to ∼100k bp downstream of the transcription start site of the long isoform of *PRDM1*.

### ATAC-seq footprint analysis

To conduct differential footprint analysis between the disease groups and cell types, we generated group-level bam files by merging the deduped bam files within each of the eight groups. A total of 47 high-quality ATAC-seq samples with FRiP>0.1 were included (MS group: 26 mTregs, control group: 21 mTregs). The differential footprint analysis was performed using HINT v0.12.3 ^102^ and TOBIAS v0.10.1 ^62^. In the analysis using HINT, we first called footprints using the command “rgt-hint footprinting --atac-seq --paired-end” with the group-specific bam files and peaks as the input files. We then identified predicted binding sites using the command “rgt-motifanalysis matching” for the 579 JASPAR (2018) core motifs for vertebrates. Finally, we identified footprints showing differential binding activity between cell types or disease groups using the command “rgt-hint differential” based on the results from the previous two steps. The same bam files and input files were used in the analysis adopting TOBIAS, in which we first ran “TOBIAS ATACorrect” to obtain bias-corrected signals and then ran “TOBIAS FootprintScores” to obtain footprint scores. Finally, we performed the differential footprint analysis with the command “TOBIAS BINDetect”.

### Single-cell RNA-seq using 10x Genomics platform

CD4^+^ T cells and CD25^hi^ CD4^+^ T cells were negatively isolated from PBMCs separately by using Easysep human CD4^+^ T cell isolation kits and EasySep Human CD4^+^CD127^low^CD25^+^ Regulatory T Cell Isolation Kit (STEMCELL Technologies), respectively. To avoid batch effects between healthy and MS samples, and to increase the numbers of Tregs to analyze, we used hashing technology (Biolegend) to pool samples in a single run of the 10x Genomics platform. Each MS sample was processed with a paired healthy control subject matched for age, sex, and ethnicity. A total of five healthy and MS sample pairs were analyzed. 100,000 cells for each cell type were subjected to Total-seq C and Hashtag antibody staining. 2 ug per 1 million cells for Total-seq C and 1 ug per 1 million cells for Hashing antibodies were used for the staining. Cells were washed three times with PBS containing 2% FBS and four hashed samples (total CD4^+^ T cells and CD25^hi^CD4^+^ T cells from each of HC and MS) were pooled into one sample. The cellular concentration was adjusted to 1,000/μL and loaded into the 10x Genomics instrument aiming to recover 10,000 cells for library preparation and sequencing. Generated libraries were then sequenced on the NovaSeq (Illumina) with a target of 50,000 reads/cell (2 × 150 paired-end reads).

### Dual omics singIe-cell analysis

#### Cell type assignment by protein surface markers

We define a latent indicator variable **z*_ik_*** to mark the assignment of a cell ***j*** to a cell type ***k*** and estimate the posterior probability of **z*_jk_*** = 1 by the stochastic expectation maximization (EM) algorithm. We assume that a normalized vector for each cell **x*_j_*** follows von Mises-Fisher (vMF) distribution ^103^ with cell type ***k***-specific mean vector **μ*_k_*** and shared concentration parameter 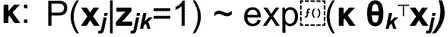. We modified the existing EM algorithm ^104^ and enforced the sparsity of the mean vector **μ** based on the prior knowledge of the cell-type-specific activity of marker proteins/genes/features. Simply, we allow **μ*gk*** to take non-zero values if and only if a feature ***g*** is a known marker for the cell type ***k***.

To improve the quality of inference, we also take advantage of “negative” marker protein labels and build an “adversarial” model for each cell type and contrast with the likelihood of the corresponding “positive” model.

● nTconv: CD3+, CD4+, CD8-, CD25-/CD127+, CD45RA+/CD45RO-
● mTconv: CD3+, CD4+, CD8-, CD25-/CD127+, CD45RA-/CD45RO+
● nTreg: CD3+, CD4+, CD8-, CD25+/CD127-, CD45RA+/CD45RO-
● mTreg: CD3+, CD4+, CD8-, CD25+/CD127-, CD45RA-/CD45RO+

To further dissect the cell types within mTreg cells, we sorted single-cell CITE-seq vectors based on the following definitions:

● Treg1: CD183+ / CD194- / CD196-
● Treg1/17: CD183+ / CD194- / CD196+
● Treg17: CD183- / CD194+ / CD196+
● Treg2: CD183- / CD194+ / CD196-

#### Batch correction and visualization of scRNA-seq data

We used the top 100 principal components of the log-transformed scRNA-seq data matrix to characterize intercellular similarity and clustering patterns across ∼45k cells and ∼15k genes. We managed to adjust discrepancies across five different batches using the batch-balancing k-nearest neighborhood method ^105^ followed by adjustment of principal component, subtracting out the mean difference between batches ^106^.

#### Single-cell differential expression analysis

We compared cell-type-specific gene expression profiles between the MS and HC subjects by estimating unbiased subject-level pseudo-bulk profiles for each gene using CoCoA-diff ^35^. CocoA-diff can improve the statistical power in case-control scRNA-seq study while adjusting for unwanted confounding effects existing across individuals. We first estimated latent factors, which may confound gene and cell-type-specific expressions with the disease labels. We established a controlled baseline for each cell derived from the MS subjects by imputing counterfactual gene expression values based on the 100 cells found in the HC cells in the top 10 PC space (BBKNN-based weighted average). Likewise, we imputed counterfactual values for the HC cells using the MS cells. More precisely, for each gene ***g*** and cell ***j***, we have observed **Y*gj*^16^** and counterfactual (imputed) **Y*gj*^(HC)^** if a cell ***j*** were derived from the MS; we have observed **Y*gj*^(HC)^** and counterfactual **Y*gj*^(MS^** if a cell ***j*** were from the HC. We then aggregate both factual (observed) and counterfactual cell profiles within each subject and cell type to estimate causal effects by comparing the average disease effect (ADE), average disease effect in the disease subject ^107^, and average disease effect in the control subject (ADC). Denoting the subject-level aggregate profiles **λ*gi*** and **λ*gi*** (for gene ***g*** and subject ***i***), we define ADE of a gene ***g*** as: ADE(**g**) = Σ*i*=1..10 log[**λ*gi*^16^** / **λ*gi*^(HC)^**] / 10, ADC(**g**) = Σ*i* in 5 HC subjects log[**λ*gi*^16^** / **λ*gi*^(HC)^**] / 5, ADD(**g**) = Σ*i* in 5 MS subjects log[**λ*gi*** / **λ*gi***] / 5. We implemented Bayesian inference methods in a C++ program that calculates gene-level statistics, including posterior mean and standard error, efficiently handling ten thousand genes and hundred thousand cells ^35^.

### Data and code availability

The accession number for the whole transcriptome sequencing data, dual omics single-cell sequencing data, and processed data reported in this paper will be provided in GEO/SRA.

## Acknowledgements

We thank patients and families, members of the Yale MS clinic; Lesley Devine and Chao Wang for technical assistance with flow cytometry; Mei Zhong and the staff of the Yale Stem Cell Center for technical assistance with bulk RNA-seq and ATAC-seq sample preparation and sequencing.

## Author contributions

Conceptualization: T.S.S., M.R.L., L.H., Y.P., M.K., and D.A.H. Recruitment of study participants: T.S.S., M.R.L., and D.A.H. Specimen collection: T.S.S., M.R.L., and H.A.S. Human primary cell experiments: T.S.S., H.A.S., and G.A.L. Mint-ChIP experiment and analysis: T.S.S., M.R.L., L.H., C.B.E. and B.E.B. Transcriptomic and eQTL analysis of ImmuNexUT data: M.O., K.F. Data analysis: T.S.S., M.R.L., L.H., and Y.P. Visualization: T.S.S., M.R.L., L.H., and Y.P. Writing review & editing: T.S.S., M.R.L., L.H., Y.P., A.M.K., M.K., and D.A.H. wrote the manuscript, with input from all co-authors. Supervision: M.K., and D.A.H.

## Supplementary Figures

**Figure S1.**
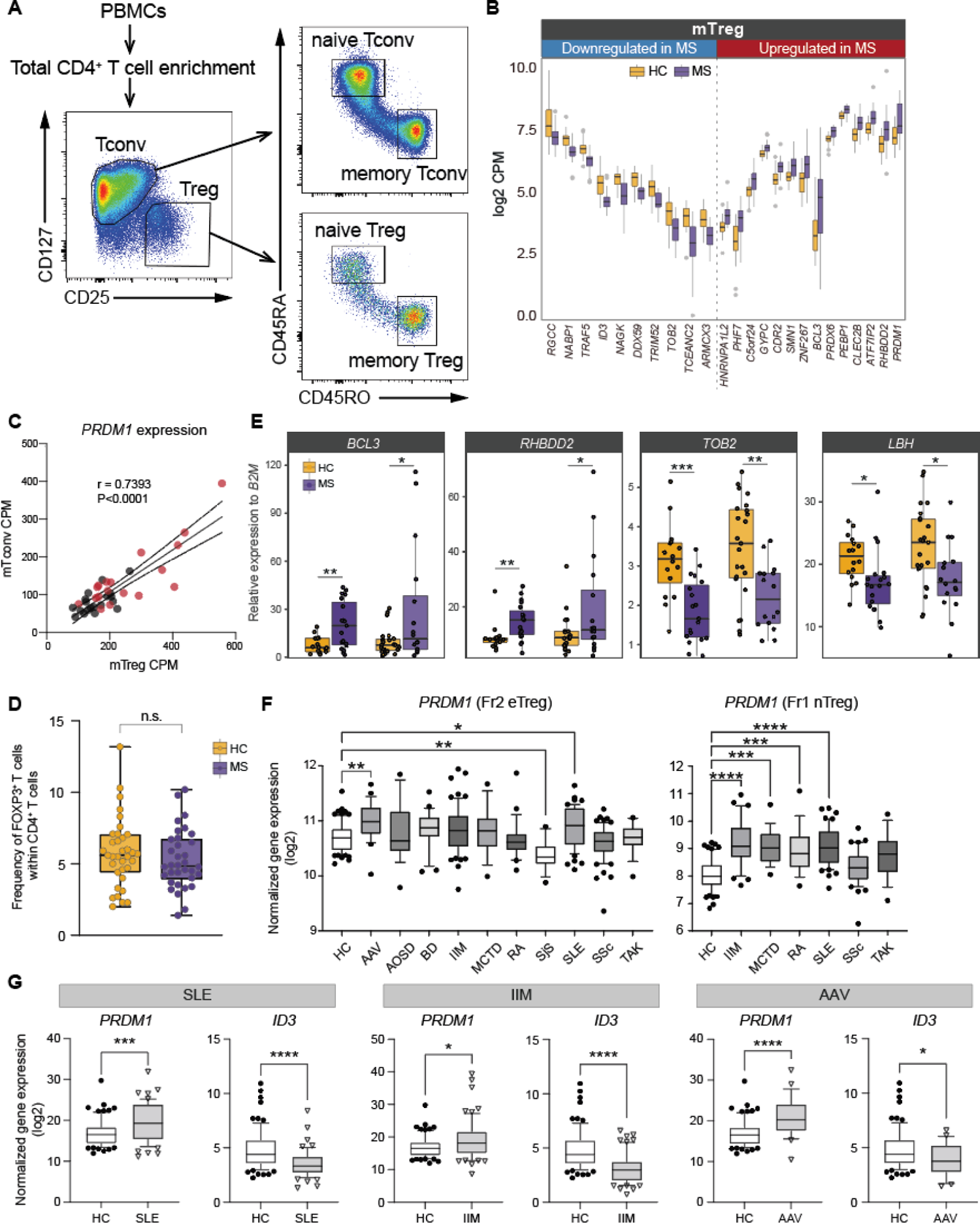
Transcriptomic analysis in mTreg and mTconv with MS patients, related to Figure 1. **(A)** Experimental workflow for isolating four major CD4^+^ T cell subpopulations by FACS. **(B)** Top individual DEGs expression at subject level. **(C)** Correlation of *PRDM1* expression in mTreg and mTconv. **(D)** Flow cytometry-based assessment of the frequency of FOXP3^+^ T cells in HC vs MS. (n=34 (HC), 35 (MS)) **(E)** qPCR validation for top four DEGs in mTreg. Data for the discovery cohort (left side) and validation cohort (right side) are shown for each HC (yellow) and MS (purple). **(F)** *PRDM1* expression in Fr2 eTreg and Fr1 nTreg across 10 and 6 autoimmune diseases are shown respectively (data were extracted from M. Ota *et al.*). **(G)** *PRDM1 and ID3* expression in Fr2 eTreg from HC, SLE, IIM, and AAV are shown respectively (data were extracted from M. Ota *et al.*) P*<0.05, P**<0.01, P***<0.001, P****<0.0001; Statistical significance computed by one way ANOVA with Dunn’s multiple comparisons tests.

**Figure S2.**
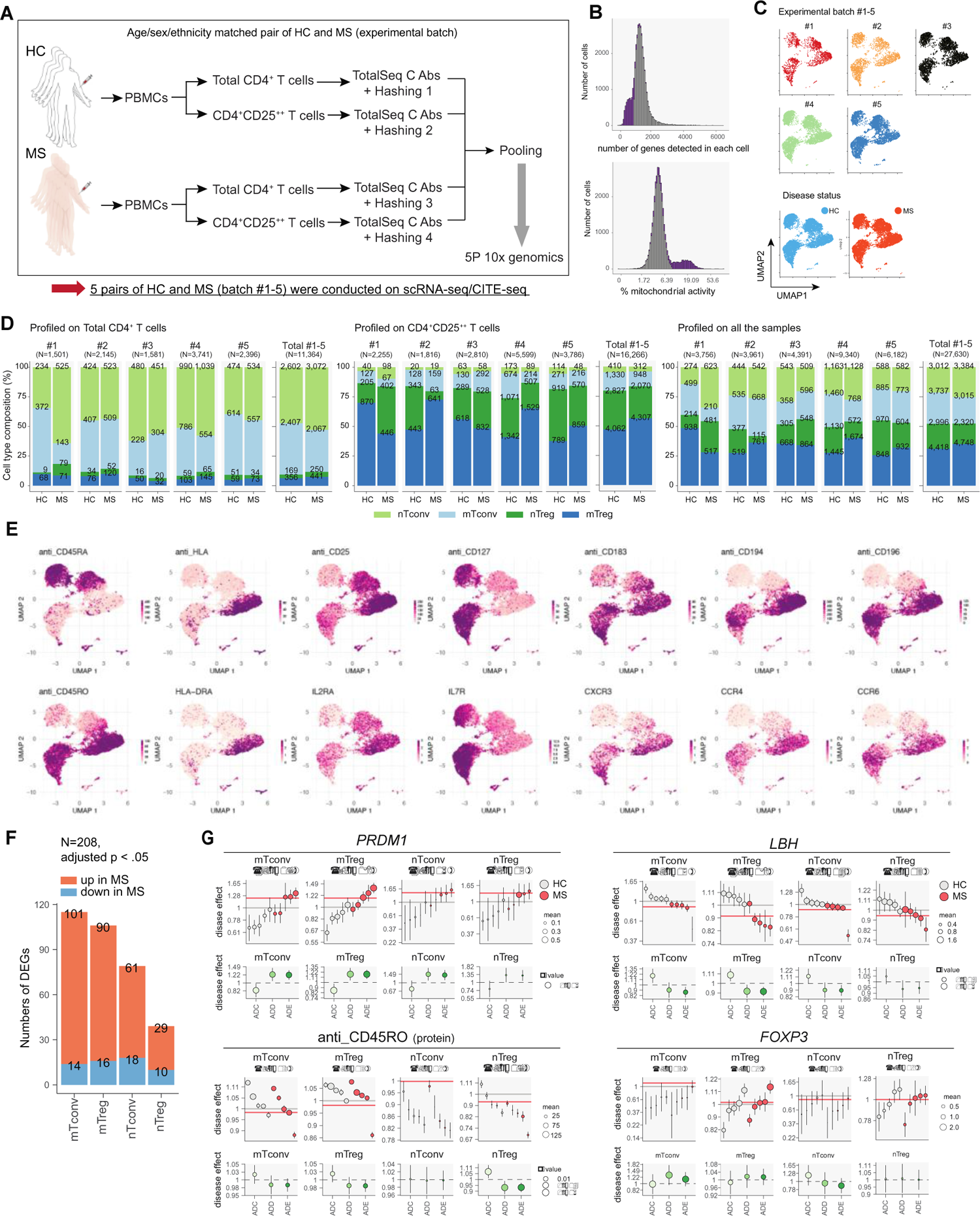
Single-cell dual omics analysis with CD4^+^ T cells in MS, related to Figure 2. **(A)** Experimental workflow for dual omics single-cell analysis of HC and MS CD4^+^ T cells. Age, sex, and ethnicity matched HC and MS subject are processed at the same time as one experimental batch. Total five experimental batches were included in this study. **(B)** Histograms showing numbers of genes detected per cell (left) and frequency of mitochondrial genes per cell (right). Cells before and after quality control selection are highlighted in purple and light gray respectively. **(C)** Gene expression UMAP of all cells color coded for experimental five batches (#1-5) (top) and disease condition (bottom). **(D)** Annotated cell numbers of CD4^+^ T cell subpopulation within total CD4^+^ T cells (left), CD4^+^CD25^++^ T cells (middle), and combined total cells (right). Cell numbers for each batch and summary of five batches from HC and MS are shown. **(E)** Representative gene and protein expressions UMAP. **(F)** Numbers of upregulated and downregulated DEGs in each CD4^+^ T cell subpopulation are shown. **(F)** Representative differential gene analysis in each CD4^+^ T cell subpopulation is depicted (See also Methods).

**Figure S3.**
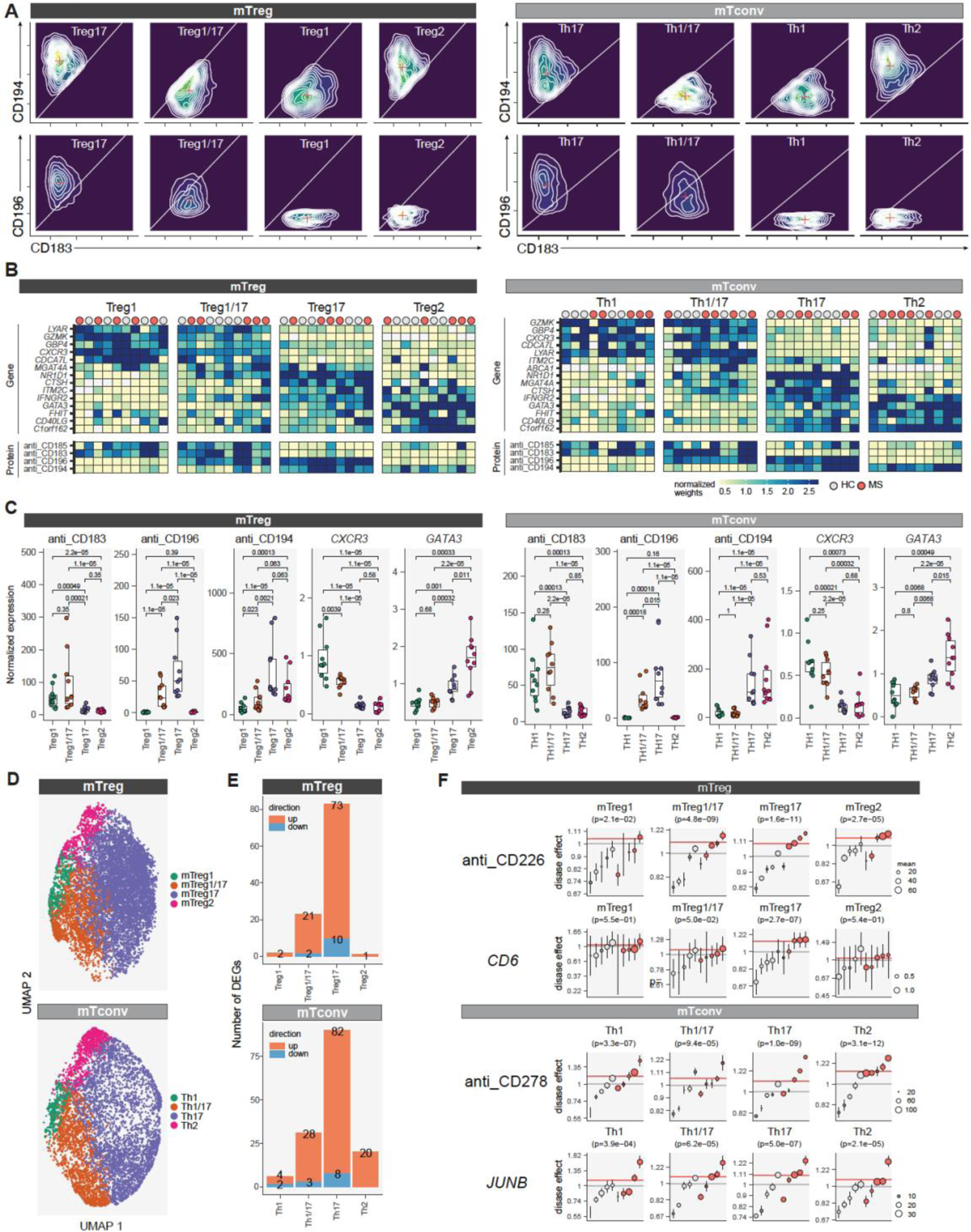
Single cell dual omics analysis with CD4^+^ T cells sub cell types in MS, related to Figure 2. **(A)** Surface protein guided mTreg and mTconv subtype annotation. CD196, CD183, and CD194 expressions were shown in four subtypes for each mTreg and mTconv. **(B)** Heatmaps showing the marker genes and proteins to define subtypes for each mTconv and mTreg at individual subject level. **(C)** Key marker expressions (CD196, CD183, CD194, *CXCR3* and *GATA3*) for each subtype in mTreg and mTconv. **(D)** UMAP based on CITE-seq based protein expressions for the subtypes in mTreg and mTconv. **(E)** Numbers of upregulated and downregulated DEGs in each subtype for mTreg (top) and mTconv (bottom). **(F)** Representative differential gene and surface protein analysis in each subtype in mTreg and mTconv are depicted (See also Methods).

**Figure S4.**
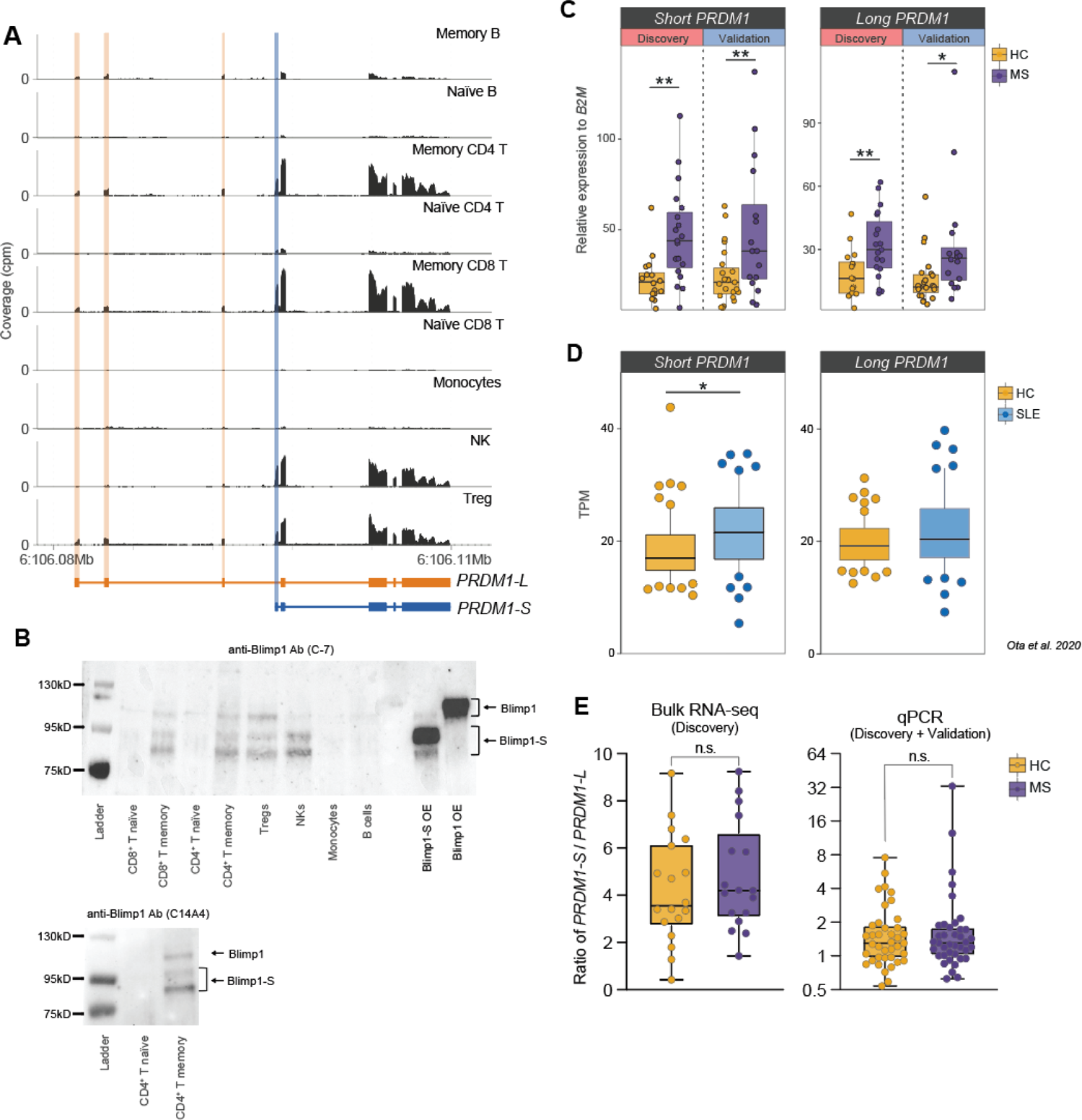
Elevated alternative short *PRDM1* isoform in MS mTreg, related to Figure 3. **(A)** Representative bulk RNA-seq coverage tracks at *PRDM1* locus from nine different immune cell types in peripheral blood. Unique exonic regions for *PRDM1-L* and *PRDM1-S* are highlighted in orange and blue respectively. **(B)** Western blot analysis of Blimp1 expression from 8 different immune cell types in peripheral blood and Blimp1-S or Blimp-1 overexpressed (OE) 293T cells by anti-Blimp1 Ab (C-7) (top) and anti-Blimp1 Ab (C14A4) (bottom). **(C)** qPCR validation of short and long *PRDM1* isoform expression between HC and MS from discovery cohort (left box plot) and validation cohort (right box plot). P*<0.05, P**<0.01; Statistical significance computed by unpaired t test. **(D)** Short and long *PRDM1* isoform expression between HC and SLE from ImmuNexUT data. **(E)** Ratio of *PRDM1-S* vs *PRDM1-L* in bulk RNA-seq and qPCR data. P*<0.05, P**<0.01; Statistical significance computed by Mann-Whitney test.

**Figure S5.**
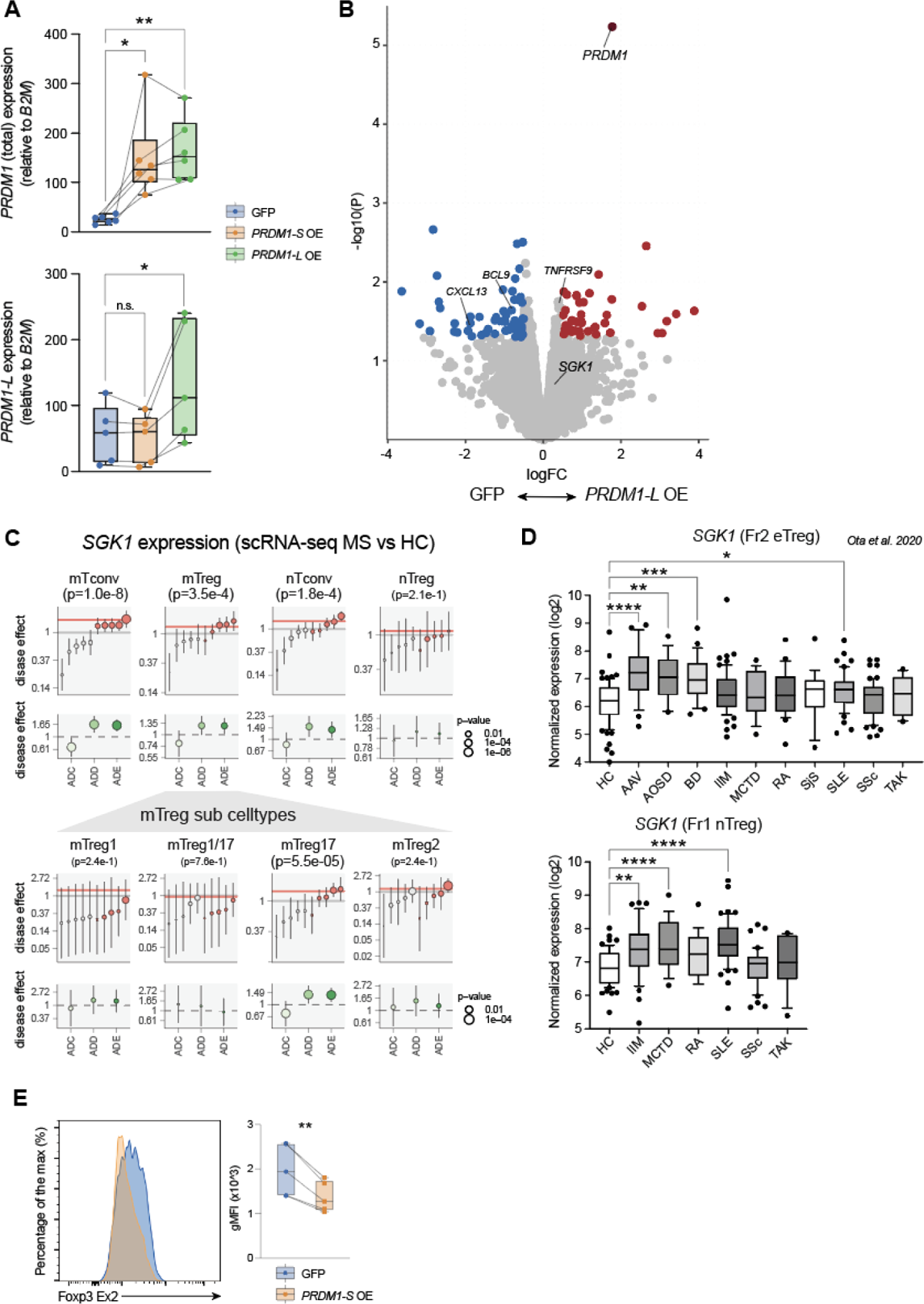
Short *PRDM1* induces *SGK1* and Treg instability, related to Figure 4. **(A)** qPCR validation of total *PRDM1* and long *PRDM1* expression by short and long *PRDM1* isoform overexpression in mTreg. P*<0.05, P**<0.01; Statistical significance computed by paired t test. **(B)** Volcano plot showing statistical significance and fold change for genes differentially expressed by long *PRDM1* overexpression in primary mTregs. **(C)** *SGK1* expression assessed by scRNA-seq in four main CD4^+^ T cell subpopulations (top) and mTreg sub cell-types (bottom). ADE: average disease effect between disease cells and matched healthy cells across all the individuals, both MS and HC. ADC: average disease effect only measured within the healthy control group. ADD: average disease effect only measured within the disease group. **(D)** *SGK1* expression in Fr2 eTreg and Fr1 nTreg across 10 and 6 autoimmune diseases respectively (data were extracted from *M. Ota et al*.). P*<0.05, P**<0.01, P***<0.001, P****<0.0001; Statistical significance computed by one way ANOVA with Dunn’s multiple comparisons tests. **(E)** Flow cytometry analysis for the specific splicing isoforms of FOXP3 containing exon 2 (FOXP3-Ex2) in primary Treg cells by overexpression of *PRDM1-S* compared to GFP control (n=5). P**<0.01; Statistical significance computed by paired t test.

**Figure S6.**
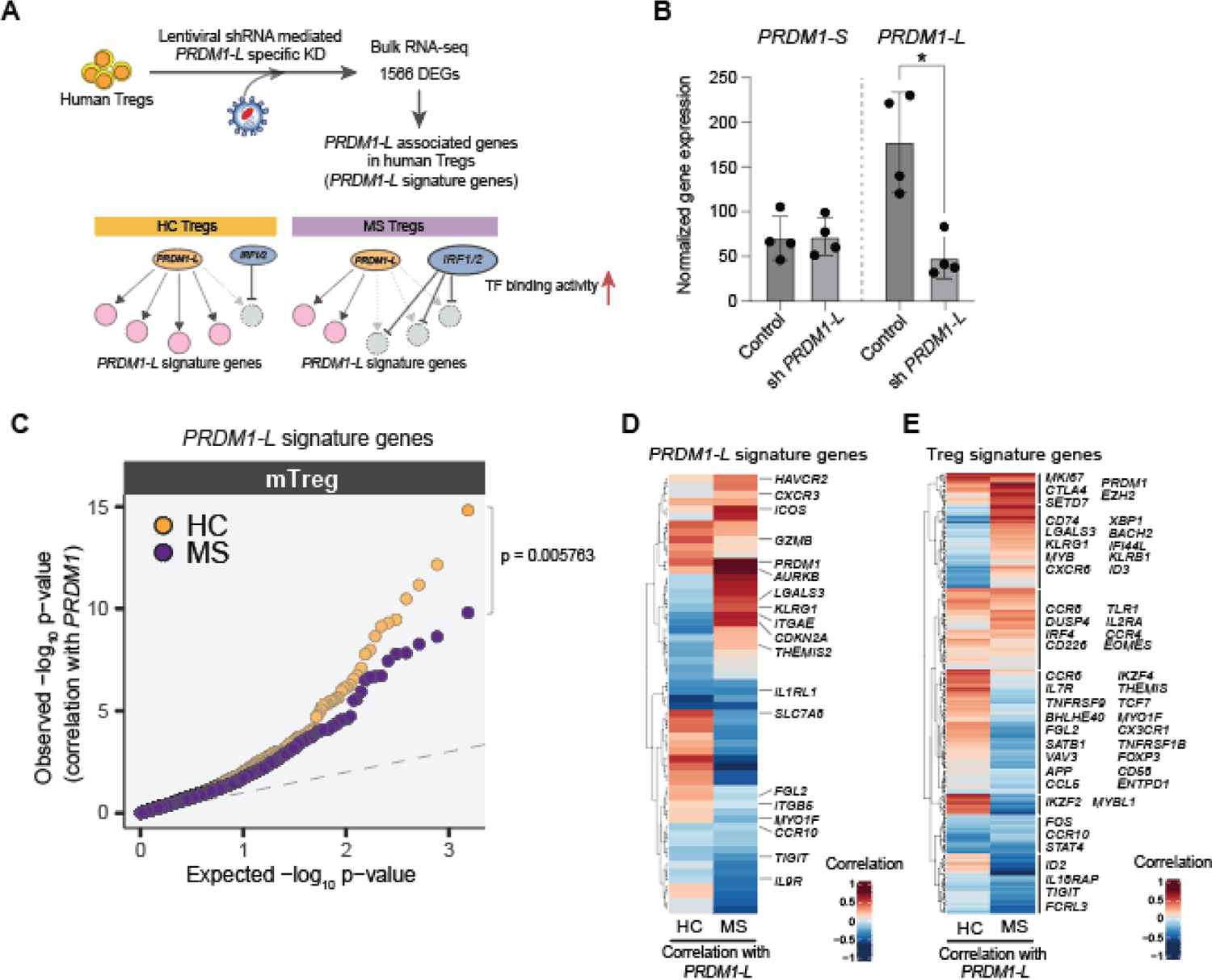
Disrupted *PRDM1-L mediated gene regulation in MS mTregs*, related to Figure 5. **(A)** Schematic of how *PRDM1-L* mediated gene regulation is disrupted by enriched TF binding of IRF1/2. **(B)** Normalized gene expression for *PRDM1-S* and *PRDM1-L* on human primary Tregs with and without *PRDM1-L* specific gene knockdown by lentiviral shRNA transduction. Lentiviral transduced GFP^+^ cells were sorted by FACS at day 5. mRNAs were isolated and bulk RNA-seq was performed. **(C)** Quantile-quantile plot showing the co-expression correlation between *PRDM1-L* signature genes with *PRDM1* at a single-cell level in mTreg for MS and HC. Correlation between *PRDM1-L* signature genes and *PRDM1* expression is stronger in HC compared to MS. Wilcoxon’s test p-value as summary values between MS vs HC: p-value = 0.005763. **(D, E)** Heatmaps depicting the Spearman’s correlation between *PRDM1-L* expression and *PRDM1-L* signature genes **(D)** and Treg signature genes **(E)**.

**Figure S7.**
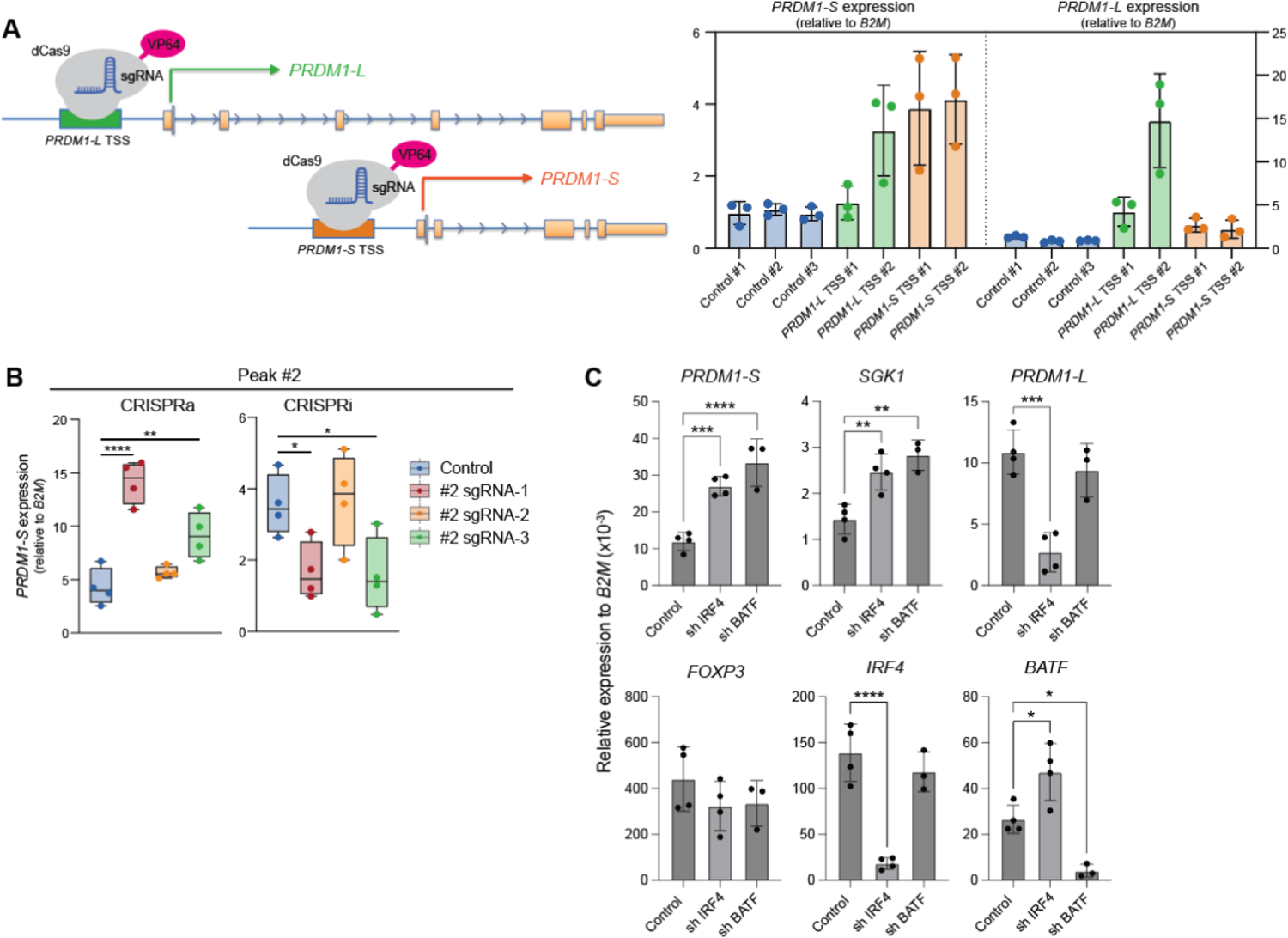
Identification of active enhancer for short *PRDM1* in human T cells, related to Figure 6. **(A)** Schematic of CRISPRa experiment for short and long *PRDM1* induction with targeting each promoter element (left) and qPCR quantification of short and long *PRDM1* expression (right). **(B)** Validation CRISPRa and CRISPRi experiment for #2 peak independent from Figure 6B (n=4). Peak #2 sgRNA-1 and -3 were validated as functional *cis*-regulatory elements for *PRDM1-S*. **(C)** Lentiviral shRNA-based gene knockdown for IRF4 and BATF in human primary Tregs. Human primary Tregs are isolated by FACS and stimulated with anti-CD3/CD28 antibodies. Lenti particles were transduced at day 1 and GFP^+^ cells were sorted by FACS at day 4-5. Expressions of each gene assessed by qPCR are shown. P*<0.05, P**<0.01, P***<0.001, P****<0.0001; Statistical significance computed by one way ANOVA with Dunn’s multiple comparisons tests.

**Figure S8.**
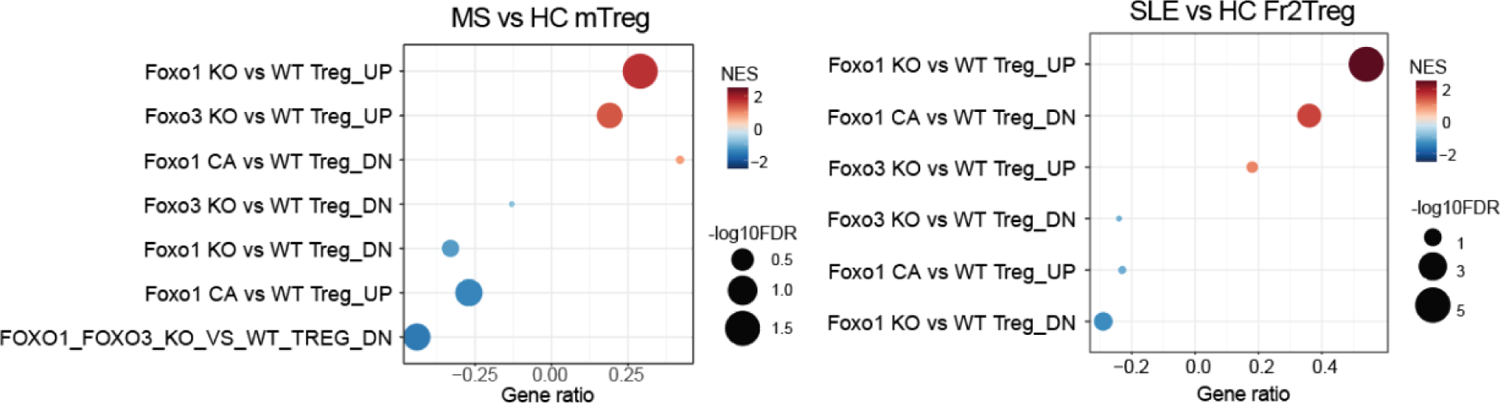
Dysfunctional Foxo1 and Foxo3 KO Treg signatures in MS and SLE mTregs. Dot plots showing gene set enrichment analysis (GSEA) for Foxo signaling on Tregs in MS (left) and SLE (right) Tregs. Gene sets generated by Foxo1/3 KO Tregs and Foxo1 constitutive active (CA) Tregs are used. NES; Normalized enrichment score.

